# Endothelin-1 from Tongue Glands Gates Sodium Taste and Intake

**DOI:** 10.64898/2026.10.28.762212

**Authors:** Li-Tzu Chen, Xinyue Xia, Ernesto R. Soto, Gary R. Ostroff, Jin Zhang

## Abstract

Sodium is an essential nutrient. To survive sodium deficiency, animals dramatically increase their appetite for salt. However, whether and how sodium depletion impacts the taste of salt to drive sodium intake is unknown. Here we show that tongue gland-derived endothelin-1 (EDN1) regulates the amiloride-sensitive, sodium-selective salt taste. Edn1 is specifically expressed in the glands embedded within the tongue. Under normal conditions, EDN1 suppresses salt taste by acting on its receptor EDNRB on the projections of salt-sensing taste neurons innervating taste buds. When internal sodium levels drop, EDN1 expression decreases, lifting this inhibition and enhancing responses to salt. Applying EDN1 directly to the tongue, using a nonionic eutectic-based organogel (Neogel) delivery technology that we establish here, dampens the activity of the salt-sensing taste neurons and reduces salt intake and preference. Conversely, genetic deletion of Ednrb in taste neurons mimics the effects of sodium deficiency, enhancing salt-sensing neuron activity and increasing salt consumption. These findings reveal a peripheral mechanism that regulates salt taste and intake in response to internal sodium levels, highlighting a direct role of the taste pathway in maintaining nutrient homeostasis.

---

Our health and survival rely on the alignment of nutrient levels and intake. Sodium is an essential nutrient, and its consumption is gated by both taste and internal state. Indeed, the taste-guided feeding response to sodium is transformed by internal sodium levels: animals normally find low concentrations of NaCl (<100mM) mildly appetitive, whereas higher concentrations become aversive (>200mM); however, salt depletion dramatically increases attraction to sodium salt in animals, including humans^1–8^. This remarkable behavioural switch towards sodium salt is crucial for restoring sodium balance and ensuring survival. While sodium deficiency is uncommon in modern diets, excessive sodium intake remains a major risk factor for cardiovascular disease—the leading cause of death worldwide—highlighting the importance of understanding how sodium intake is regulated.

How do animals and humans adjust sodium consumption according to internal sodium levels? Nearly a century ago, Richter and colleagues pioneered studies on sodium deficiency in rats^9,10^ and proposed that heightened salt preference is not learned through experience but instead arises from “chemical changes in taste mechanisms in the oral cavity, giving rise to an enhanced salt discrimination”^10^.

Here, we uncover a peripheral mechanism that gates salt taste in response to sodium deficiency. We show that the endothelin-1 (EDN1) from tongue glands suppresses salt-sensing neuron activity and salt intake through EDNRB. Salt depletion decreases EDN1 expression, lifting this inhibition and enhancing salt taste and consumption.

## Salt depletion transforms the taste of salt

To test how internal sodium levels impact salt consumption, we monitored blood and urinary sodium and serum aldosterone levels in mice before and after sodium depletion. To induce sodium depletion, animals were injected intraperitoneally (i.p.) with furosemide, a diuretic drug, and given ad libitum water and sodium-free food to replace lost fluids and electrolytes while preventing sodium intake^11^. This treatment effectively decreased blood and urine sodium levels^12,13^, accompanied by changes in serum aldosterone (Fig. 1a, Extended Data Fig. 1a–b).

**Figure 1.**
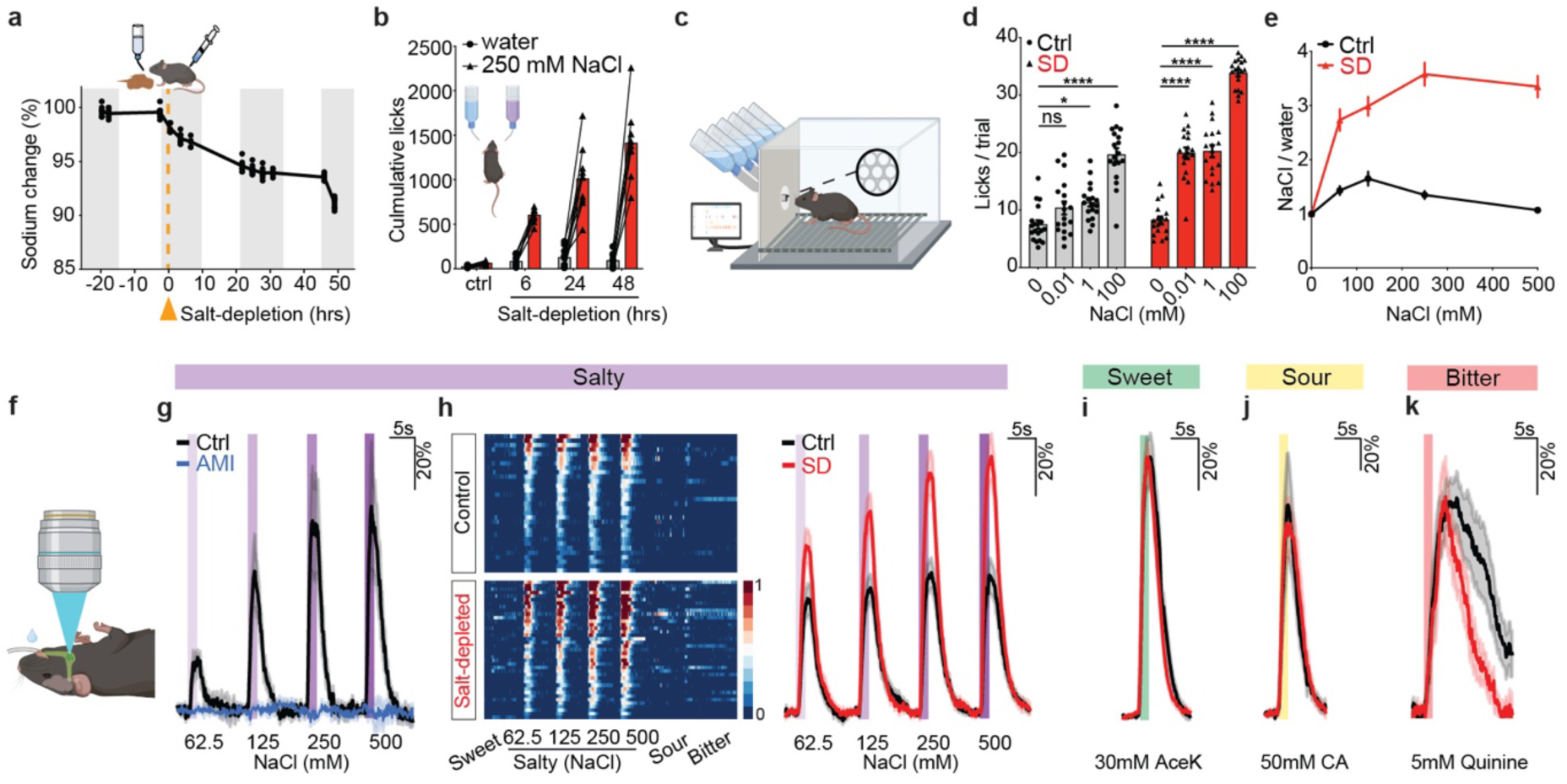
Sodium depletion induces a sensitized salt taste and enhanced activity of salt-sensing taste neurons. **a.** Blood sodium levels decrease gradually in salt-depleted (SD) animals. Dark cycle (grey). Black line represents mean. Animals were injected with furosemide intraperitoneally at time 0 (orange) and provided water and sodium-free food (*n* = 10). **b.** Two-bottle taste preference test with water and 250mM NaCl. Cumulative licks for water (grey) versus NaCl (red) were shown for animals that received a mock injection (Ctrl), or 6, 24, and 48 h post SD (*n* = 10). Paired t-test, *P* = 0.0002 (Ctrl), *P* < 0.0001 (SD 6, 24, 48 h) **c.** Custom-engineered gustometer with the layout of the drinking spout enlarged (see Methods). **d.** Licks per 5s trial to 0.01, 1, and 100mM NaCl in controls (Ctrl, grey) and 48 h SD (red). Values are mean ± s.e.m., *n* = 18, One-way ANOVA, Dunnett’s multiple comparisons test. Ctrl, *P* = 0.0152 (1mM), *P* < 0.0001 (100mM); SD, *P* < 0.0001 (0.01, 1, and 100mM). **e.** Lick ratio of NaCl to water in Ctrl and SD animals. Values are mean±s.e.m., *n* = 10, Two-way ANOVA, Šídák’s multiple comparisons test, *P* < 0.0001. **f.** Diagram illustrates optical access to the geniculate ganglion. Tastant solutions were delivered to the tongue. Responses are normalized to the mean sweet-evoked response of sweet-selective neurons (**g-k**). **g.** Salt-tuned neuronal responses are eliminated by amiloride (AMI). Normalized average responses of 8 salt-tuned neurons from 5 Ctrl mice before (black) and after AMI treatment (blue). **h-k.** Calcium imaging of taste ganglion neurons in Ctrl and SD animals. *n* = 451 neurons in 19 mice (Ctrl); *n* = 720 in 25 mice (SD); **h.** Left, heatmaps depicting fluorescence traces from NaCl-selective neurons. Each row represents the average activity of a single cell in two trials. Stimuli are listed at the bottom: sweet (30mM Acesulfame potassium, AceK), salt (62.5, 125, 250, and 500mM NaCl), sour (50mM citric acid), and bitter (5mM quinine). *n* = 31 (Ctrl), *n* = 39 (SD). Right, average responses to NaCl in Ctrl (black) and SD (red), normalized to the mean sweet response in each animal. Shaded areas (s.e.m.). Bars represent 2s stimulation windows. **i-k.** Average responses of sweet-(**i**, Ctrl, *n* = 68, SD, *n* = 106), sour-(**j**, Ctrl, *n* = 16, SD, *n* = 22), and bitter-(**k**, Ctrl, *n* = 15, SD, *n* = 11) selective neurons, normalized to the mean sweet response in each animal. Unnormalized traces are shown in Extended Data Fig. 2. Schematics in **a,b,c,f** were created using BioRender (http://biorender.com).

We tested animals for salt preference using a two-bottle preference test at 6, 24, and 48 hours after furosemide injection. As internal sodium levels progressively decreased, sodium consumption increased (Fig. 1a–b). Importantly, salt depletion not only increased salt preference, but also significantly enhanced taste sensitivity to NaCl, determined using our custom gustometer (see Methods, Fig. 1c). We tested control and salt-depleted animals using NaCl concentrations spanning 0.01–100 mM. While control animals showed a detection threshold of ∼1 mM NaCl, salt-depleted animals detected and showed significantly increased licking at 0.01 mM NaCl (Fig. 1d, Extended Data Fig. 1c). These enhanced taste responses to NaCl after salt depletion were evident at higher concentrations (62.5–500 mM, Fig.1e) and were progressively enhanced at 6, 24, and 48 hours after salt depletion (Extended Data Fig. 1d). By contrast, taste responses to sweet, bitter, or non-sodium salts such as potassium, calcium, and magnesium salts were unaffected (Extended Data Fig. 1e-i).

## Peripheral salt-tuned neurons are sensitized after salt depletion

In mice, sodium is detected by ENaCα/β/γ-expressing taste receptor cells (TRCs), an amiloride-sensitive salt-sensing pathway that promotes sodium attraction and consumption^1,14,15^, whereas higher concentrations of NaCl (>200 mM), as well as other salts such as potassium and magnesium, additionally recruit amiloride-insensitive bitter- and sour-sensing pathways, triggering aversion^2^. The vastly enhanced preference and sensitivity to NaCl in salt-depleted animals suggest sensitized salt taste that can provide an elevated drive for sodium intake, which we hypothesized may result from enhanced activity of salt-sensing peripheral taste neurons. We directly tested this hypothesis by imaging taste-evoked neuronal activity in geniculate ganglion (GG) neurons that innervate fungiform papillae of the anterior tongue (Fig. 1f), where the amiloride-sensitive sodium-specific TRCs primarily reside^1,16–19^. Most GG neurons are singly tuned to one taste modality^20–23^, offering a key advantage for distinguishing functionally distinct neuronal populations and evaluating population-specific changes following salt depletion with single-cell resolution.

To record and compare taste-evoked neuronal activity between control and salt-depleted animals, we expressed GCaMP in all GG taste neurons by crossing *P2rx3-Cre*^20^ with *FLEx-GCaMP6s*^24^ lines. Because salt-tuned GG neurons innervate the amiloride-sensitive, sodium-selective TRCs, we confirmed that their NaCl-evoked responses were amiloride-sensitive, whereas bitter- and/or sour-tuned neurons responding only to higher concentrations of NaCl were amiloride-insensitive (Extended Data Fig. 2a–f). Importantly, we found a dramatic increase in the responses of amiloride sensitive salt-tuned GG neurons to the same NaCl titrations in salt-depleted animals compared with controls (Fig. 1h). In contrast, the activity of neurons tuned to other taste modalities was comparable to control conditions (Fig. 1i–k, Extended Data Fig. 2j–r). This heightened activity in salt-tuned GG neurons may contribute to enhanced taste detection and increased salt intake.

## EDN1–EDNRB as a candidate ligand–receptor pair regulating salt taste

To identify potential mechanisms and gene candidates that directly modulate peripheral taste responses, we used two strategies: first, we cross-referenced 207 genes specifically expressed in salt-tuned GG neurons^20^ with 45 genes in the Gene Ontology (GO) term for sodium ion homeostasis. This yielded a single candidate: *Ednrb*, which encodes endothelin receptor B (EDNRB) and is 23.8-fold enriched in the salt-sensing cluster compared to other taste-tuned clusters^20^ (Extended Data Fig. 3a–b). Second, hypothesizing that a relevant gene would exhibit altered expression after salt depletion, we curated a list of 14 genes known to mediate sodium detection or salt balance and measured their expression in peripheral taste tissues by quantitative RT-PCR (RT-qPCR). These candidates include genes encoding the sodium salt taste receptors, ENaCα,β,γ^1,14^, and fluid homeostasis effectors in the Renin, Angiotensin, and Aldosterone pathways^25,26^. Among them, only *Edn1*, which encodes endothelin-1, showed a significant, progressive decrease at 6, 24, and 48 hours following salt depletion (Extended Data Fig. 4a).

Endothelin peptides (EDN1, EDN2, and EDN3) bind to EDNRA and EDNRB receptors and play key roles in fluid homeostasis in the cardiovascular and renal systems^27–29^. Given that *Ednrb* is enriched in salt-sensing taste ganglion neurons and *Edn1* expression in the tongue decreases with internal sodium loss, we hypothesized that this ligand-receptor pair modulates salt-responsive neuronal activity. Because GG neurons innervate the tongue and transmit taste signals to the rostral nucleus of the solitary tract (rNST)^30–32^, we examined expression of endothelin family members in the tongue, GG, and the rNST using RT-qPCR. In contrast to *Edn1* and *Ednrb*, other endothelin family members showed low or unchanged expression across these regions (Extended Data Fig. 4).

## *Edn1* is expressed in tongue glands

To identify the specific tongue cell types expressing *Edn1* and to compare gene expression differences between control and salt-depleted animals, we performed scRNA-seq on the tongue epithelium, enriching for taste bud populations by micro-dissecting taste papillae from both the anterior (fungiform) and posterior tongue (circumvallate and foliate, Fig. 2a). This approach effectively resolved diverse epithelial and taste bud cell types based on their distinct gene expression profiles (Fig. 2b–d, Extended Data Fig. 3c–f). In addition to taste-related populations such as type I (*Entpd2*^+^)^33^, type II (*Trpm5*^+^)^34,35^, type III (*Pkd2l1*^+^) cells^36–38^, the proliferating stem cells (*Mki67*^+^), and lingual progenitors (*Lgr5*^+^)^39^, our scRNA-seq analysis generated a comprehensive map revealing previously uncharted populations in the tongue epithelium (Fig. 2b–d, Extended Data Fig. 3).

**Figure 2.**
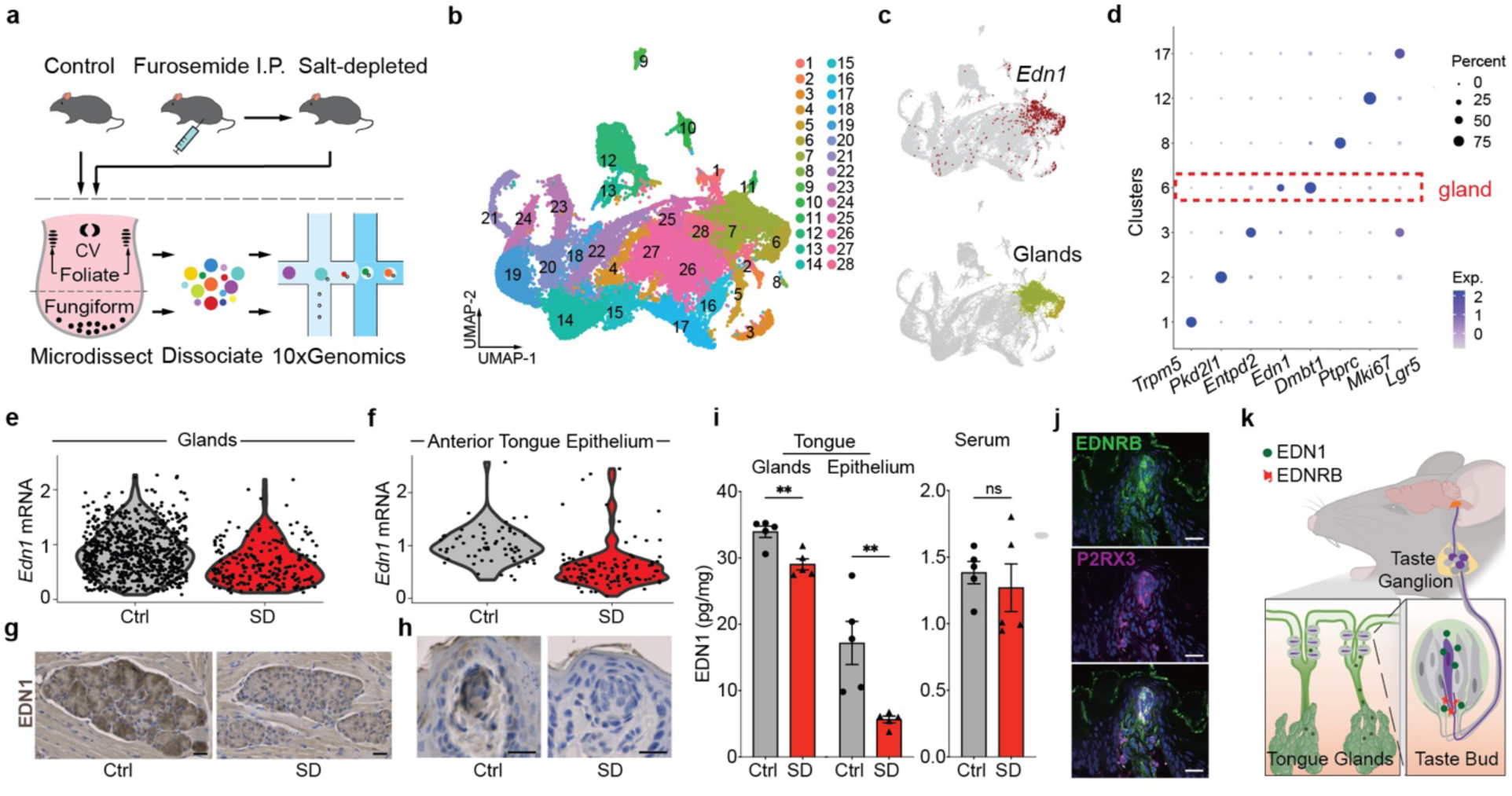
Endothelin 1 is expressed in the tongue glands, and its expression decreases after salt depletion. **a.** Tongue epithelia of control and salt-depleted animals were isolated, areas containing taste buds, including circumvallate (CV), foliate, and fungiform papillae, were micro-dissected, and cells were dissociated for 10x genomics scRNA-seq analysis. **b.** UMAP showing 28 segregated clusters of tongue epithelial cells. **c.** *Edn1* is primarily expressed in the glands. Top, cells expressing *Edn1* (normalized expression >1) are colored red; Bottom, clusters 6 (lime) and 7 (green) label the gland populations; both overlaid on the UMAP (grey) shown in **b**. **d.** Dot plot showing marker gene expression. *Edn1* is mainly expressed in cluster 6, which co-expresses *Dmbt1*. *Trpm5* marks type II taste cells, *Pkd2l1* type III sour-sensing cells, *Entpd2* type I glia-like cells, *Ptprc* putative leucocytes, *Mki67* putative proliferating stem cells, and *Lgr5* lingual progenitors. **e-f.** *Edn1* gene expression is significantly decreased in 24 h salt-depleted animals compared with controls in both glands (**e**; Ctrl, *n* = 768 cells; SD, *n* = 219; *P* = 9.435 × 10⁻¹⁴) and anterior tongue epithelium (**f**; Ctrl, n = 64; SD, n = 106; P = 5.225 × 10⁻¹⁰), obtained by micro-dissection of fungiform papillae. **g-h.** Haematoxylin-eosin and EDN1 staining of von Ebner’s glands (**g**) and a representative fungiform taste bud (**h**) in Ctrl and SD animals (*n* = 4). **i.** EDN1 protein levels are reduced in salt-depleted tongue epithelium and glands (embedded in the tongue muscle) but are unchanged in serum compared with controls. *n* = 5, unpaired t-test, *P* = 0.0035 (gland); *P* = 0.0081 (epithelium); *P* = 0.5796 (serum). **j.** Immunostaining for EDNRB (green) and P2RX3 (magenta) at the projections of ganglion neurons innervating a representative taste bud (*n* = 4 mice). **k.** Cartoon illustrating EDN1 released from the glands embedded in the tongue, binding to EDNRB on the projections of salt-sensing neurons innervating taste buds. All scale bars, 50µm. Schematics in **k** were created using BioRender (http://biorender.com).

We discovered that *Edn1* is almost exclusively expressed in gland populations (clusters 6 and 7) and was not detected in taste bud cells (e.g., types I, II, III) from either the anterior or posterior tongue (Fig. 2c–d). The gland populations are marked by *Dmbt1* (Deleted in Malignant Brain Tumors 1), whose gene products have been detected in the ducts of tongue-embedded glands, including the von Ebner’s glands (VEG), and are released into the fluid bathing taste buds^40,41^. Consistent with this finding, immunostaining and H&E images revealed that both EDN1 and DMBT1 are localised to the VEG (Extended Data Fig. 5a–b).

## EDN1 levels decrease after salt depletion

Consistent with *Edn1* RT-qPCR analysis (Extended Data Fig. 4), our scRNA-seq data showed diminished *Edn1* mRNA in salt-depleted animals in gland and anterior tongue epithelial populations (Fig. 2e–f). Although *Edn1* mRNA is almost exclusively expressed in tongue glands, EDN1 protein is present in both glands and taste buds, with levels significantly decreased at both sites after salt depletion, despite unchanged gland and taste-bud size (Fig. 2g– i, Extended Data Fig. 5d–e). In contrast, EDN1 levels in blood serum did not change after salt depletion (Fig. 2i), supporting a local source of EDN1 in the minor glands embedded in the tongue.

*Ednrb,* which encodes an EDN1 receptor, is highly and specifically expressed in salt-sensing GG neurons^20^ (Extended Data Fig. 3–4). To localize EDNRB within these neurons, we performed double labelling and found it enriched at ganglion neuron terminals marked by P2RX3–a purinergic receptor essential for receiving and transmitting taste signals^42–44^ (Fig. 2j, Extended Data Fig. 5f). These results indicate that EDN1, produced by tongue glands and present in taste buds, potentially signals through EDNRB at salt-tuned GG neuronal terminals to regulate salt taste activity (Fig. 2k).

## EDN1 administration decreases salt intake and dampens neuronal activity

Since decreased EDN1 levels correlated with enhanced salt taste responses following salt depletion, we hypothesized that EDN1–EDNRB signalling inhibits salt taste responsiveness and salt intake. We therefore tested whether EDN1 administration would decrease taste responses to NaCl in salt-depleted animals. As EDN1 levels peaked in the fungiform taste buds 30 min post-injection (Fig. 3a–b), we tested animals’ behavioural responses to NaCl 30 min after EDN1 i.p. injection. EDN1-injected animals displayed significantly attenuated taste responses to different concentrations of NaCl (Fig. 3c), indicating that EDN1 suppresses salt taste responses. In contrast, injecting EDN2 or EDN3 did not change taste behaviour to NaCl (Extended Data Fig. 7a–c).

**Figure 3.**
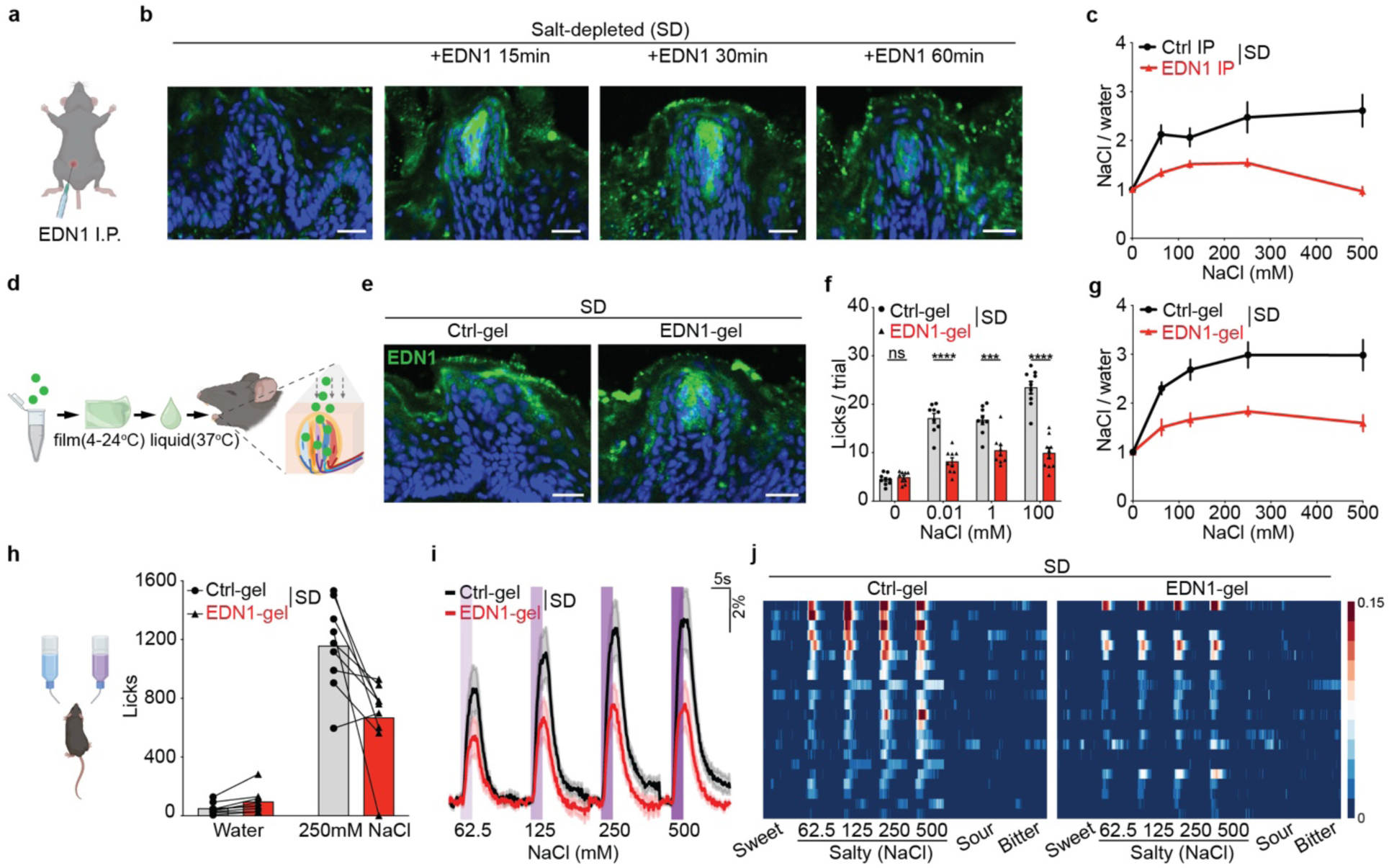
EDN1 application suppresses salt taste responses, reduces salt intake, and dampens activity of salt-sensing neurons. **a.** EDN1 intraperitoneal (i.p.) injection. **b.** EDN1 (green) is lowly expressed in taste buds of salt-depleted animals (left) but enriched at 15, 30, and 60 min after i.p. injection. DAPI (blue). A representative fungiform taste bud is shown for each condition (*n* = 3 mice). **c.** EDN1 i.p. injections significantly decrease animals’ lick ratio of NaCl to water. *n* = 10, mean ± s.e.m., two-way ANOVA, Šídák’s multiple comparisons test, *P* = 0.0009. **d.** Cartoon illustrates the Neogel delivery platform. The gel adopts a solid form at room temperature and liquefies at body temperature, facilitating cargo delivery. **e.** EDN1 (green) immunostaining in a representative fungiform taste bud 30 min after application of empty Neogel (Ctrl-gel) or Neogel containing EDN1 (EDN1-gel). *n* = 3 mice. **f.** EDN1-gel attenuates salt taste sensitivity. Licks per 5 s trial to 0.01, 1, and 100 mM NaCl in salt-depleted animals treated with Ctrl-gel (grey) and EDN1-gel (red). Values are mean ± s.e.m.; n = 9; *P* < 0.0001 (0.01 and 100 mM), *P* = 0.0002 (1 mM). **g.** EDN1-gel application decreases the animals’ lick ratio of NaCl to water. *n* = 10, mean ± s.e.m., two-way ANOVA, Šídák’s multiple comparisons test, *P* = 0.002. **h.** Taste preference test where mice chose between water and 250mM NaCl. Bars represent cumulative licks to 250mM NaCl in salt-depleted animals after receiving Ctrl-gel (grey) or EDN1-gel (red). *n* = 9, paired t-test, *P* = 0.009 (250mM NaCl). **i.** Calcium responses of salt-selective taste ganglion neurons in the same salt-depleted animals after receiving Ctrl-gel (black) or EDN1-gel (red). Traces represent average responses (dF/F); shaded areas indicate s.e.; purple bars represent 2s stimulation windows of 62.5, 125, 250, and 500mM NaCl. *n* = 22 salt-selective neurons out of 248 recorded (Extended Data Fig. 8). **j.** Heatmaps show responses of salt-selective taste ganglion neurons in **h** to different taste stimuli. Each row represents the average activity of a single cell in two trials. Stimuli are listed at the bottom: sweet (30mM Acesulfame potassium, AceK), salt (62.5, 125, 250, and 500mM NaCl), sour (50mM citric acid), and bitter (5mM quinine). All scale bars, 50µm. Schematics in **a,d,g** were created using BioRender (http://biorender.com).

To test whether EDN1 directly alters taste detection, we developed a highly specific, non-invasive topical delivery system targeting the tongue epithelium (Fig. 3d). After screening 15 different matrices, we identified one that had no effect on taste behaviour when applied alone (Extended Data Fig. 6). This non-ionic eutectic-based organogel (Neogel) forms a solid film at room temperature that adheres to the tongue and liquefies at body temperature, releasing its payload to the tongue epithelium, with signal detected in taste buds by anti-EDN1 when delivering EDN1 (Fig. 3e) and by anti-FITC when delivering FITC-conjugated EDN1 (Extended Data Fig. 7f). Using Neogel to deliver EDN1 to the tongues of salt-depleted animals significantly reduced taste sensitivity across NaCl concentrations of 0.01–100 mM, reduced licking to NaCl across 62.5–500 mM in the gustometer, and reduced NaCl preference in two-bottle preference assays (Fig. 3f–h). In contrast, EDN1 did not impact food or water intake nor change sweet or bitter tastes (Extended Data Fig. 7).

To investigate whether EDN1 administration suppresses salt neuron activity, we imaged salt-depleted *P2rx3-Cre, FLEx-GCaMP6s* mice, which express a calcium indicator in all taste neurons^20^. We first applied solvent-containing Neogel (Ctrl-gel) to the tongue of a salt-depleted animal and conducted calcium imaging to capture the baseline activity of GG neurons. We then administered EDN1-containing Neogel (EDN1-gel), and imaged GG neurons of the same animal 30 min after application. EDN1-gel treatment directly attenuated the activity of amiloride-sensitive salt-tuned GG neurons, but did not significantly impact neurons tuned to other tastes or the amiloride-insensitive bitter- and sour-tuned neurons that respond to concentrated NaCl (Fig. 3i–j, Extended Data Fig. 8). Together, systemic and local applications of EDN1 revealed that EDN1 suppresses salt-selective taste responses, inhibits salt-sensing taste neuron activity, and decreases salt intake.

## EDNRB in taste neurons is required for EDN1 to suppress the taste of salt

Given that reductions in EDN1 levels drive increased consumption and taste responses to NaCl, we predicted that knocking out its receptor in taste neurons should lead to increased salt taste sensitivity, heightened neuronal responses, and elevated salt consumption independent of salt depletion. To test this prediction, we generated conditional knockout animals that specifically delete *Ednrb* in taste ganglion neurons (*P2rx3-Cre*^20^; *Ednrb ^f/f^* ^45^, *Ednrb cKO,* Fig. 4a, Extended Data Fig. 9). Indeed, *Ednrb cKO* animals showed increased sensitivity to NaCl (Fig. 4b), recapitulating what was observed in salt-depleted animals (Fig. 1d). These *cKO* mice also exhibited significantly greater salt intake and higher salt-to-water ratios across 8 hours of continuous monitoring in two-bottle preference assays compared with littermate controls (*P2rx3-Cre*; *Ednrb^+/f^*, *Ednrb het,* Fig. 4c–e).

**Figure 4.**
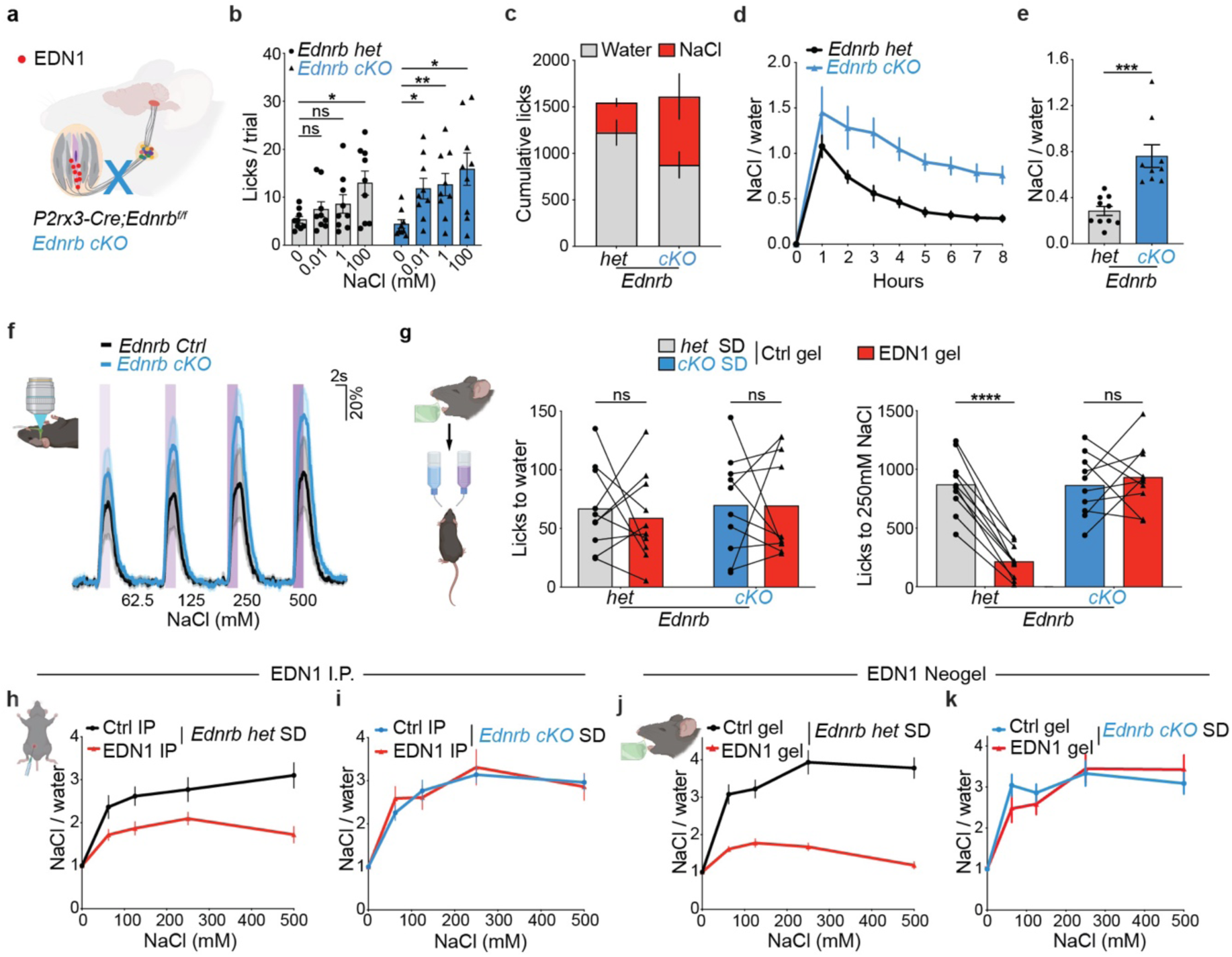
*Ednrb* loss sensitizes salt taste, mimicking salt depletion and blocking EDN1 effects. **a.** Cartoon illustrating the *Ednrb* conditional knockout (cKO) mice. **b.** Animals’ licks per 5s trial to 0.01, 1, and 100mM NaCl in *Ednrb het* (grey) and *cKO* (blue) animals. *n* = 9 each. One-way ANOVA, Dunnett’s multiple comparisons test, *Ednrb het*, *P* = 0.5037 (0.1mM), *P* = 0.4299 (1mM), *P* = 0.0285 (100mM); *Ednrb cKO*, *P* = 0.0143 (0.1mM), *P* = 0.0058 (1mM), *P* = 0.0188 (100mM). **c-e.** Taste preference test in which mice chose between water and 500 mM NaCl. **(c)** Cumulative licks recorded over 8 h in *P2rx3-Cre; Ednrb^+/f^* (*het*, *n* = 10) and *P2rx3-Cre; Ednrb^f/f^* (*cKO*, *n* = 9) mice. **(d)** NaCl-to-water lick ratios over 8 h. Two-way ANOVA with Šídák’s multiple-comparisons test, *P* = 0.0017. (**e**) NaCl-to-water lick ratios, unpaired t-test, *P* = 0.0002. **f.** Calcium responses of salt-selective taste ganglion neurons in *Ednrb Ctrl* (grey) and *cKO* (blue) animals. Traces represent average responses of salt-selective neurons normalized to the mean sweet-evoked response of sweet-selective neurons. Shaded areas (s.e.m.). Purple bars represent 2s stimulation windows of 62.5, 125, 250, and 500mM NaCl. n=14 neurons (*Ednrb Ctrl*), n=12 (*Ednrb cKO*). **g.** Taste preference test in which salt-depleted mice chose between water and 250 mM NaCl. Bars show cumulative licks for water (left) or 250 mM NaCl (right) in *Ednrb* het (gray) and cKO (blue) mice after application of empty Neogel (Ctrl-gel) or EDN1-containing Neogel (EDN1-gel). Paired *t*-test, for 250 mM NaCl: *P* < 0.0001 (*Ednrb* het) and *P* = 0.5759 (cKO). **h-i.** EDN1 i.p. injections significantly decrease the NaCl-to-water lick ratio in salt-depleted *Ednrb het* but not in *Ednrb cKO* animals. *n* = 10 each, Two-way ANOVA, Šídák’s multiple comparisons test, *P* = 0.0040 (**h,** *het*); *P* = 0.8598 (**i**, *cKO*). **j-k.** Tongue application of Neogel containing EDN1 reduces taste responses to NaCl in salt-depleted *Ednrb het* but not in *cKO* animals. *n* = 10, Two-way ANOVA, Šídák’s multiple comparisons test, *P* < 0.0001 (**j**, *het*), *P* = 0.7834 (**k**, *cKO*). Schematics in **a,f,g,h,j** were created using BioRender (http://biorender.com).

To examine whether the enhanced taste sensitivity stems from increased neuronal activity, we expressed GCaMP6s in the ganglion neurons of *Ednrb cKO (P2rx3-Cre*; *Ednrb ^f/f^*; *GCaMP6s*) and controls. Salt-tuned neurons in *Ednrb cKO* animals showed heightened NaCl-evoked activity compared with control littermates when stimulated with the same NaCl titrations (Fig. 4f), substantiating enhanced NaCl-evoked responses in non-depleted *Ednrb cKO* animals.

If EDN1 acts through the EDNRB receptor expressed in GG neurons to suppress salt intake, then exogenous EDN1 should have no effect in animals lacking *Ednrb* in these neurons. Indeed, EDN1 attenuated salt preference, salt taste responses, and consumption in salt-depleted *Ednrb het*, but not in *Ednrb cKO* animals, in two-bottle preference assays (Fig. 4g) and following either i.p. injection (Fig. 4h–i) or Neogel delivery in gustometer analysis (Fig. 4j–k). These results demonstrate that EDNRB expressed in peripheral taste neurons is required for EDN1 to suppress the taste of salt.

## Discussion

Our findings reveal a key peripheral mechanism regulating sodium taste and intake during sodium deficiency. After sodium depletion, animals and humans immediately start ingesting sodium when it reaches the tongue–well before it can be absorbed or detected by internal sensors^5–9^ (Fig. 1). This rapid behavioural switch led us to hypothesize that the taste system itself is adaptively tuned to promote sodium intake under deficiency. While several studies have focused mainly on central circuits^4,26^, recordings of the chorda tympani, which contains peripheral fibers of GG neurons innervating the anterior tongue, showed no change or suppression of salt responses during sodium depletion^46–50^, leaving the role of the peripheral taste system unclear. Salt taste responses are also particularly complex, with amiloride-sensitive salt-sensing TRCs primarily residing in fungiform papillae of the anterior tongue, while amiloride-insensitive bitter- and sour-sensing pathways can respond to concentrated NaCl^1,2^. Our single-neuron calcium imaging distinguishes changes from functionally distinct GG populations, which can be obscured by aggregate nerve recordings. Here, we identify EDN1 produced by tongue glands as a local regulator of the amiloride-sensitive salt-sensing pathway through EDNRB on salt-tuned taste neurons. Crucially, this inhibition is lifted after sodium depletion, enhancing salt taste and increasing salt intake. These findings reveal an active and dynamic role for the taste system in adapting sodium intake to internal sodium state and contributing to sodium homeostasis.

The endothelin family has been extensively studied for its critical roles in regulating systemic blood pressure and sodium balance^27,29,51^. While the function of EDN1 in promoting sodium excretion in the kidney is well established, its role in regulating sodium intake has not been explored. Notably, high sodium levels increase EDN1 expression in the kidney, activating EDN1–EDNRB signalling to facilitate sodium excretion^27,29^. Our findings identify a homeostatic function of EDN1–EDNRB signalling in the taste pathway, whereby reduced EDN1 following sodium depletion enhances salt taste and intake. Together, these functions position EDN1–EDNRB signalling as a key regulator of sodium homeostasis through both sodium excretion and intake.

As *Edn1* levels were highest in control animals and progressively decreased after salt depletion (Extended Data Fig. 4a), we sought to determine whether this modulation is driven by oral sodium detection or restoration of internal sodium levels. We treated animals with furosemide to induce sodium depletion and then administered NaCl either by oral licking or i.p. injection. Interestingly, just 5 minutes after oral NaCl administration, *Edn1* expression in the tongue rose significantly, suggesting rapid regulation of local *Edn1* expression by oral sodium detection. In contrast, administering the same amount of NaCl via i.p. had no effect on *Edn1* levels even after 30 min (Extended Data Fig. 10). Consistent with our findings, previous studies have shown that oral exposure to NaCl triggers an immediate decrease in pre-LC^PDYN^ “salt appetite” neuron activity, whereas intragastric infusion does not^52^. Together, these results suggest that oral sodium detection provides a rapid signal before systemic sodium balance is restored and raise the possibility that dynamic *Edn1* expression in the tongue links sodium detection to adaptive salt intake.

Our finding that *Edn1* is expressed in tongue-embedded glands is particularly intriguing. These glands have long been proposed to secrete peptides that influence taste perception^53–55^, although their functions have remained elusive. Their ducts can connect directly to taste buds (Extended Data Fig. 5a–b), illustrating a potential anatomical route through which gland-derived factors may influence taste. Although the anatomical route by which gland-derived EDN1 reaches the anterior tongue remains unresolved, identifying EDN1 as a gland-derived modulator of salt taste supports this proposed role and opens new avenues to investigate other gland-secreted factors that may regulate taste and feeding behaviour.

To examine the function of EDN1, we developed a Neogel-based platform to deliver it locally to the tongue surface. This non-invasive delivery platform enables targeted application of peptides and small molecules to modulate taste at the site of detection. In addition to providing a strategy for suppressing salt intake, it could be adapted to investigate other taste modalities or potentially help curb excessive consumption of highly palatable foods, such as sweets.

In summary, we have identified a peripheral mechanism that directly regulates sodium detection and ingestion in response to internal sodium state. The ability to enhance sodium detection and increase its intake during sodium depletion is a key sensory adaptation to sodium deficiency. Understanding how internal nutrient states shape taste perception is increasingly important, as disruption of this coordination contributes to major health burdens—including hypertension, obesity, and diabetes. In the future, it will be of great interest to elucidate how the taste system integrates other internal signals to gate nutrient intake and whether this mechanism becomes dysregulated in disease.

## Methods

### Animals

All procedures were carried out in accordance with the US National Institutes of Health (NIH) guidelines for the care and use of laboratory animals and were approved by the University of Massachusetts Chan Medical School’s Institutional Animal Care and Use Committee. Experiments were carried out using adult mice aged between 6 weeks and 8 months. In behavioural assays, animals of the same age and sex composition were used in the control and experimental groups. For transgenic animals, littermates were used as controls. *P2rx3-Cre* was crossed with the Cre-dependent GCaMP6s reporter line (JAX028866) for calcium imaging of the geniculate ganglion. The *Ednrb* conditional knockout animals were generated by breeding *P2rx3-Cre* with *Ednrb^f/f^* (JAX011080).

### Salt depletion and Physiological Measurements

To induce salt depletion, mice received intraperitoneal injections of furosemide (50 mg/kg; Sigma, F4381) and were maintained on a salt-deficient diet (Inotiv, TD90228). To prevent coprophagic sodium reuptake, cages were changed three times per hour during the first 3 h post-injection, and again at 6, 24, and 48 h (Extended Data Fig. 1a).

To assess food and water intake throughout salt depletion and Neogel treatment, animals were singly housed, and food and water intake were measured daily during the dark cycle. Food intake was recorded as the amount of food consumed per animal per day, and water intake was recorded as the volume of water consumed per animal per day.

Blood was collected from the tail vein, and urine was collected every 3 h during the dark cycle for analysis. Serum sodium levels were measured using a sodium assay kit (Sigma, MAK247), and aldosterone levels were measured using an ELISA kit (Enzo Life Sciences, ADI-901-173), according to the manufacturers’ instructions.

### Custom-engineered gustometer behavioural analysis

To accurately assess taste detection while minimizing extraneous cues, we engineered a custom behavioural chamber based on the widely used gustometer^34,56^ but incorporated three key modifications: (1) all taste solutions were delivered through separate channels converging into a single spout, eliminating potential bias or learned associations with specific spouts; (2) the spout was thoroughly rinsed between trials to remove any residual sensory cues; and (3) each channel delivered 1–2 μL of liquid per lick, enabling high-resolution analysis of taste detection. The chamber was built on an extra-wide modular test chamber (Med Associates) inside a sound-attenuating cubicle, using a combination of 3D-printed and laser-cut components. The spout contained seven internal channels—six connected to different concentrations of taste solutions, and one central channel connected to a vacuum line for efficient removal of residual liquid during cleaning. An electronic lick sensor was attached to the spout, and spout access was regulated by a mechanical shutter. The entire system was automated and controlled using Arduino and MATLAB. Each trial lasted 5 seconds and was initiated by the animal’s first lick, with licks automatically recorded.

Mice were acclimated to the custom gustometer and trained to lick water from the spouts 2 days prior to tests. For behavioural testing of salt responses, the water bottle was removed for 6–8 h prior to the tests to prevent differences in water consumption before testing and to standardize the animals’ internal state. Animals were then provided water trials in the gustometer, and testing was initiated only when water licking did not exceed 20 licks per 5-s window. This screening procedure was used to minimize the potential contribution of thirst to the behavioural measurements. For attractive tastants (sucrose and AceK), animals were food-deprived for 20 h and water-deprived for 4-6 h prior to the tests. For behavior towards bitter tastants, mice were water-deprived for 24 h prior to the tests. All animals were placed individually into the gustometer chamber and presented with one tastant per trial. Each 15-30 min test session consisted of 18 trials: the first 6 trials presented water and increasing concentrations of a tastant to allow animals to sample each stimulus, followed by 12 randomized trials. The number of licks, lick latency, and inter-lick interval were automatically recorded for each trial.

### Two-bottle preference assays

Two-bottle preference assays were conducted in the animals’ home cages using a custom-designed 3D-printed scaffold that held two bottles. Each bottle was connected to an electronic lick sensor, allowing continuous and automated recording of licking throughout the experiment. To acclimate the animals to the setup, mice were given access to two bottles filled with water for two consecutive days prior to testing. On the day of the experiment, water was removed for 1 h before the assay to prevent immediate consumption before the test and to standardize the animals’ internal state. During the test, one bottle contained water and the other contained either 500 mM or 250 mM NaCl. Water and salt intake were continuously monitored for 1h (Fig.1b, 3h, and 4g) or 8 h (Fig. 4c–e).

### Calcium imaging and analysis

Animal surgery in preparation for calcium imaging was described previously^22^. Anesthetized mouse was put at a supine position with its head fixed to a metal bar with dental cement. The tracheotomy was performed to secure the airway and prevent choking during oral tastant delivery.

Imaging data were obtained using the Evolve EMCCD camera (16-micron pixel, 512x512, Teledyne Photometrics). All images were acquired at 10Hz, 100ms exposure, with 10x magnification. Tastants were delivered (5–10 mL per min) using silicone tubing positioned approximately 8 mm inside the oral cavity, dorsal to the tongue. Tastants dissolved in artificial saliva were delivered for 2s each in serial order, interspersed with 10s or 13s of artificial saliva rinse. Images were acquired during continuous stimulus presentation. The concentrations of tastants used were: 30 mM AceK, 5 mM quinine hydrochloride, 62.5 mM to 500mM NaCl, 50 mM citric acid. Because there are very few umami-responding neurons in the mouse geniculate ganglion^14–16^, umami tastants were not used.

To identify amiloride-sensitive salt neurons, 10 μM amiloride was added to all tastant solutions and the artificial saliva rinse. The sequential taste stimulation protocol was performed twice in the same mice, first without amiloride and then with amiloride, using the same order of tastant presentation in both conditions.

Imaging data were analyzed using custom software implemented in MATLAB as described previously^20–22,57^. To identify neurons, maps of peak activity (maximal pixel intensity over mean pixel intensity) were median filtered, threshold gated, and separated by watershed segmentation to create candidate regions of interest representing active neurons, which were reviewed manually to identify ROIs for all active neurons. Fluorescence traces for each region of interest were normalized to neighborhood fluorescence intensity (defined as the average intensity within a two-cell radius distance of each cell, excluding all other defined ROIs). Each ganglion was imaged three times. For experiments involving amiloride, each ganglion was imaged twice in the absence of amiloride and twice in the presence of amiloride to reduce the total surgery time. Images from individual trials were first concatenated and then motion-corrected using Template Matching (https://sites.imagej.net/Template_Matching/), followed by analysis using custom software implemented in MATLAB, as described previously^20–22,57^. Neurons that responded consistently in at least 2 trials were included in the analysis. Calcium transients were scored as fluorescence excursions of > 6-fold above noise, calculated as the median average deviation of the baseline 2s preceding each tastant stimulus. We also visually scored cells by directly observing the aligned image data displayed as a relative fluorescence movie and the putative cells’ fluorescence time series. To ensure that signals originated from a single neuron, and not from closely adjacent cells or out-of-focus fluorescence from deeper cell layers, we examined the correlation of pixels in the neighborhood of each ROI for each calcium transient, discarding from analysis any neuron contaminated by outside signals^20–22,57^.

### Quantitative RT-PCR

Tissue samples of circumvallate papilla, foliate, fungiform, geniculate ganglion, and rostral nucleus of the solitary tract (rNST) were collected and homogenized in TRIzol (Sigma). Total mRNA was isolated following the manufacturer’s instructions. cDNA was prepared using SuperScript II Reverse Transcriptase (ThermoFisher). Quantitative PCR was performed on QuantStudio3 (ThermoFisher) using KAPA SYBR FAST kit (KAPA biosystems). Analyzed mRNA levels were normalized to *Gapdh*.

### Cell isolation and single-cell RNA-sequencing

Tongues were extracted from euthanized mice and injected with a dispase/collagenase (Roche) enzyme cocktail (4mg/mL in HBSS). After a 30-min incubation at 37 °C, lingual epithelia were carefully peeled off from the muscle layer as previously described. Circumvallate and foliate papillae from the posterior tongues of 4 mice for each treatment condition were combined in one tube. In parallel, ∼1 mm² of epithelium from the tongue tip containing fungiform papillae from the same 4 mice was combined in a separate tube, dissected, and minced under a microdissection microscope. The two tissue preparations were processed separately but in parallel: each was incubated in 0.25% Trypsin-EDTA (Gibco) for 15 min at 37 °C, triturated, and passed through a 40-μm filter (pluriSelect) to obtain single cells. Cells were centrifuged at 500g for 5min at 4°C and resuspended in ice-cold HBSS containing 0.04% BSA. Cell densities were quantified with a hemocytometer and diluted to 400-1,000 cells/μL. A suspension of ∼10,000 single cells per sample was loaded onto the Chromium Next GEM chip G (10x Genomics) per the manufacturer’s protocol. Chromium Single Cell 3′ Gel Bead and Library Kit v3.1 and the Dual index kit TT set A (10x Genomics) were used to prepare sequencing libraries following the manufacturer’s protocol. Libraries were sequenced on Illumina Nova 6,000 sequencers (Pair-end 150bp).

### Sequencing analysis

Sequenced reads were aligned to the mouse reference genome (mm10), and gene expression matrices were generated using Cell Ranger (10x Genomics). Single-cell RNA-seq data were analyzed in R (v4.3.3) using Seurat v4^58^ (https://satijalab.org/seurat/). Custom R scripts were developed following standard Seurat workflows. For quality control, genes were retained if expressed in more than three cells, and cells were included if they expressed more than 1,000 unique genes and had less than 10% of reads mapping to mitochondrial genes. Gene expression was normalized by total expression per cell, scaled by a factor of 10,000, and log transformed. Datasets from circumvallate-foliate papillae and fungiform papillae from each treatment condition were first merged, while preserving cellular identities associated with each papilla type. The resulting datasets from control, 24 h, and 48 h sodium-depleted samples were then integrated using reciprocal PCA, with the control group as the reference. To identify cell types, the top 2,000 most variable genes were selected and scaled. Dimensionality reduction and clustering were visualized using UMAP.

### Immunohistochemistry and haematoxylin-eosin staining

Immunohistochemistry on paraffin sections was performed using tongue tissue fixed in 10% formalin for 24 h, dehydrated, and embedded in paraffin. Sections (5 μm) were cut, deparaffinized, and rehydrated. Haematoxylin-eosin staining was performed by Applied Pathology Systems. For EDN1 immunohistochemical staining, antigen retrieval was performed using a 1× Tris-based unmasking solution (Vector Laboratories, H-3301). Sections were incubated with anti-EDN1 antibody (Abcam, ab117757, 1:300), followed by SignalStain® Boost IHC Detection Reagent (Cell Signaling Technology, HRP Rabbit, 8114P) for 30 min at room temperature. Detection was performed using the SignalStain® DAB Substrate Kit (Cell Signaling Technology, 8059P). Sections were then counterstained with hematoxylin (Sigma, GHS132) and blued with bluing reagent (Epredia, 7301). After dehydration, slides were mounted with SignalStain® Mounting Medium II (Cell Signaling Technology, 84583) and imaged using a Zeiss Axio Scan.Z1 slide scanner. The areas of taste buds and von Ebner’s glands were quantified. For each image, an equal number of taste buds was randomly selected for quantification, independent of taste bud size, morphology, or staining intensity, to minimize selection bias.

### Immunofluorescence staining

Animals were euthanized and perfused intracardially with 4% paraformaldehyde solution. Tongues were excised and placed in 30% sucrose solution overnight at 4°C for cryoprotection. Tissue was embedded in OCT compound and sectioned at 16 μm thickness on a cryostat (Leica Biosystems). Sections were washed in PBS with 0.1% Triton X-100, blocked with 5% donkey serum, incubated with primary antibodies overnight at 4°C, and labelled with fluorescently tagged secondary antibodies for 3 h at room temperature. Primary antibodies used were anti-DMBT1 (ThermoFisher, PA5-115313, 1:300), anti-EDN1 (Abcam, ab117757, 1:300), anti-EDNRB (Abcam, ab117529, 1:300), anti-FITC (Abcam, ab19224, 1:100), and anti-P2RX3 (Sigma, AB5896, 1:300). A W1-Yokogawa spinning disk Nikon confocal microscope was used for fluorescence imaging.

### Endothelin-1 ELISA assay

Tongues were isolated and injected with 4mg/mL dispase/collagenase (Roche) in HBSS. After 30 min of incubation at 37 °C, the tongue epithelium, von Ebner’s glands underneath the circumvallate papillae, and the tongue muscle were dissected and isolated. Tissue samples were homogenized in protein extraction buffer containing 100 mM Tris pH 7.4, 150 mM NaCl, 1 mM EGTA, 1 mM EDTA, 1% Triton X-100, and 0.5% sodium deoxycholate, and incubated at 4 °C for 2 h with a gentle shake. Samples were then centrifuged at 13,000 g for 20 min, and the supernatant was collected for ELISA (Sigma, RAB1852) following the manufacturer’s instructions.

### The Neogel tongue delivery platform

To prepare the Neogel, 27 μL glycofurol and 23 μL lauric acid were mixed and heated at 50 °C for 10 min, and the 50μL mixture was added to a 10 x 10 x 5 mm cryomold (Tissue-Tek) and incubated at 4 °C for 30 min before use. For Neogel carrying EDN1, 12.46 μg lyophilized EDN1 was first dissolved in 27 μL of glycofurol (Sigma, T3396) and mixed with 23 μL of lauric acid (TCI L0011). The 50μL mixture was vortexed and added to the cryomold as described above, and 0.1 cm^2^ of the Neogel containing EDN1 was delivered to the tongue surface of an animal. For intraperitoneal injection, 0.1 mg/kg EDN1 (Sigma, E7764), EDN2, or EDN3 (synthesized from GenScript) dissolved in deoxygenated water to preserve endothelin function by preventing oxidation^29^. Animal behavior was tested in our custom gustometer 30 min after injection.

### Oral salt stimulation

For oral salt stimulation, salt-depleted animals were each placed in the gustometer to complete 50 licks of 1 M NaCl and were euthanized at 5- or 30-min post-stimulation. For NaCl administered i.p., salt-depleted animals each received a 50 μL injection of 1 M NaCl and were euthanized 5- or 30-min post-injection. Tongue epithelia were isolated and processed for mRNA extraction, cDNA synthesis, and quantitative PCR.

### Statistics

Statistical methods used include Dunnett’s multiple comparisons test and Šídák’s multiple comparisons test and are indicated for all figures. All the statistical tests are two-tailed. Analyses were performed in R, Python, and GraphPad Prism 9. Data are presented as mean ± s.e.m. No statistical methods were used to predetermine sample size, and investigators were not blinded to group allocation. No method of randomization was used to determine how animals were allocated to experimental groups.

### Inclusion and ethics

We support an all-inclusive, diverse, and equitable conduct of research.

## Data availability

All data supporting the findings of this study are available from the corresponding author (J.Z.).

## Code availability

Custom code is available from the corresponding author (J.Z.).

## Acknowledgements

We are especially grateful to Charles Zuker for his generous support during the early stage of this study and for his continued guidance and insightful feedback. We thank past and present members of the Zhang lab for their helpful discussions, experimental assistance, and technical support. We are indebted to Norman Kennedy for his expert guidance on immunohistochemistry staining. We appreciate Miguel Villavicencio’s guidance and help on constructing the custom gustometer. We also thank Victor Ambros, Roger Davis, Michael Francis, Paul Greer, Craig Peterson, Nick Ryba, and Andrew Tapper for their invaluable comments and discussions. This work was supported by startup funds from UMass Chan to J.Z., a Whitehall Foundation Research Grant to J.Z., and the Mello Scholar Award to L.C.

## Author contributions

J.Z., L.C., and X.X. designed the study, conducted experiments, and analyzed the data. E.S. and G.O. developed the Neogel formulation. J.Z. wrote the paper.

## Materials & Correspondence

Correspondence to

## Tables

**Table 1. Top 10 gene ontology terms enriched in top 100 marker genes of each tongue epithelial cluster.**

---

**Extended data Fig. 1.**
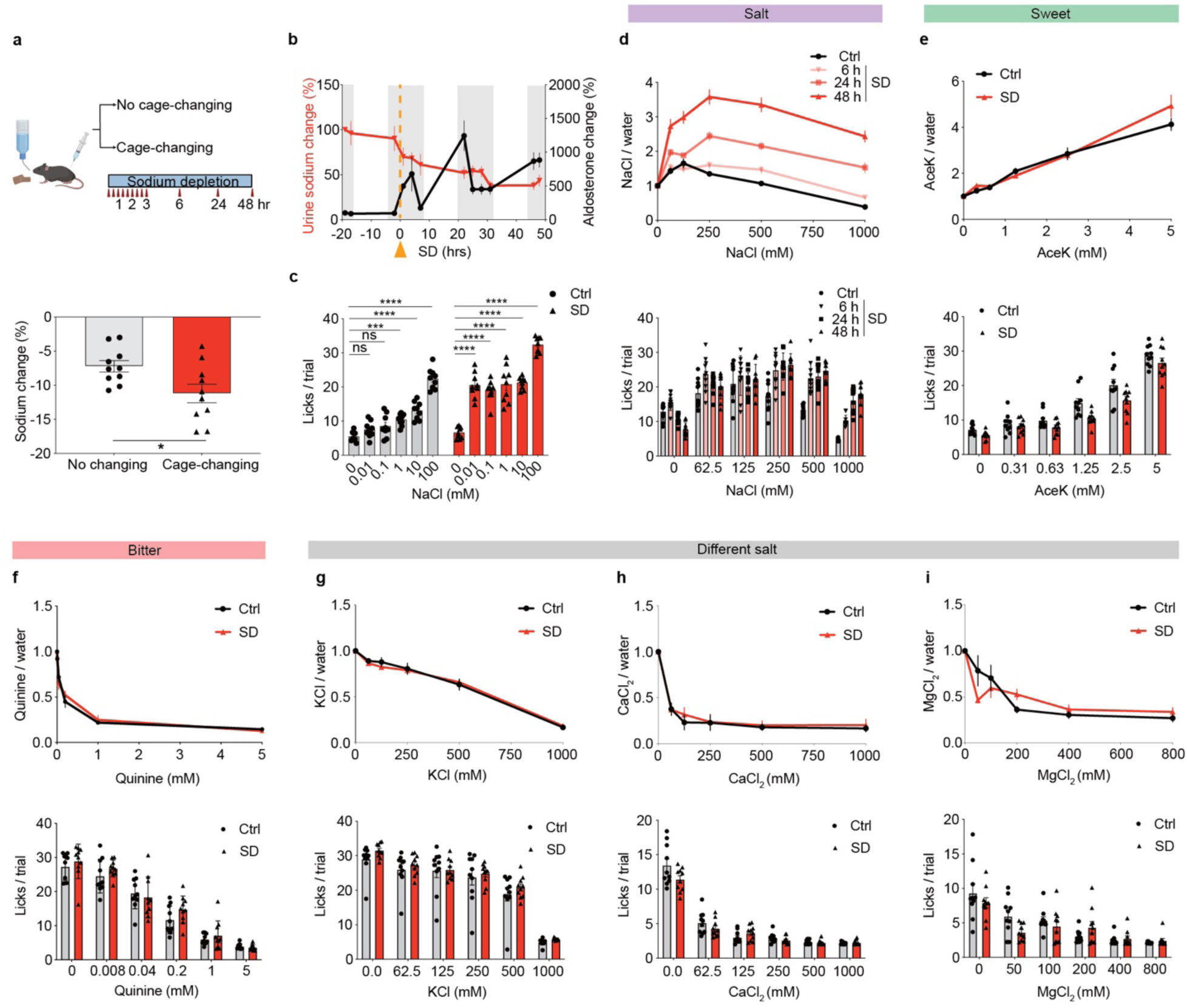
Salt depletion induces a specific and gradual increase in taste responses to sodium salt. **a.** Cage-changing after sodium depletion resulted in a significantly larger decrease in serum sodium levels. Top, sodium depletion without cage-changing and with cage-changing three times per hour in the first 3h and at 6, 24, and 48 h. Bottom, percentage decrease in sodium levels 48 h after sodium depletion relative to controls, *n* = 10, unpaired t-test, *P* = 0.0220. **b.** Changes (%) in urine sodium (red) and blood aldosterone (black) following sodium depletion (SD). Mean ± s.e.m., Dark and light cycles are shown in gray and white bars, respectively. Animals received intraperitoneal furosemide at time 0 (orange) and were provided water and sodium-free food (*n* = 10). **c.** Licks per 5 s trial to 0.01, 0.1, 1, 10, and 100 mM NaCl in controls (Ctrl, grey) and 48 h SD (red). Values are mean ± s.e.m.; *n* = 9 (responses to 0.01, 1, and 100 mM from the same animals are also shown in Fig. 1d). One-way ANOVA with Dunnett’s multiple comparisons test. Ctrl, *P* = 0.0005 (1 mM), *P* < 0.0001 (10 and 100 mM); SD, *P* < 0.0001 (all comparisons). **d.** Lick ratio of NaCl to water in control (Ctrl) and salt-depleted (SD) animals 6, 24, 48 h after furosemide injection. Bottom, licks per 5s trial to water, 62.5, 125, 250, 500, and 1,000 mM NaCl. *n* = 10, *P* = 0.0484 (6 h SD); *P* < 0.0001 (24 and 48 h SD). **c-h,** Two-way ANOVA, Šídák’s multiple comparisons test. **e.** Lick ratio of Acesulfame Potassium (AceK) to Ctrl and SD animals 48 h after furosemide injection. Bottom, licks per 5s trial to water, 0.31, 0.63, 1.25, 2.5, 5mM AceK. *n* = 9, *P* = 0.1701. **f.** Lick ratio of quinine to water in Ctrl and SD animals 48 h after furosemide injection. Bottom, licks per 5s trial to water, 0.01, 0.05, 0.2, 1, 5mM quinine. *n* = 10, *P* = 0.8366. **g.** Lick ratio of KCl to water in Ctrl and SD animals 48 h after furosemide injection. Bottom, licks per 5s trial to water, 62.5, 125, 250, 500, and 1,000mM KCl. *n* = 10, *P* = 0.7986. **h.** Lick ratio of CaCl_2_ to water in Ctrl and SD animals 48 h after furosemide injection. Bottom, licks per 5-sec trial to water, 62.5, 125, 250, 500, and 1,000mM CaCl_2_. *n* = 10, *P* = 0.0912. **i.** Lick ratio of MgCl_2_ to water in Ctrl and SD animals 48 h after furosemide injection. Bottom, licks per 5-sec trial to water, 50, 100, 200, 400, and 800mM MgCl_2_. *n* = 10, *P* = 0.7666. Schematics in **a** were created using BioRender (http://biorender.com).

**Extended data Fig. 2.**
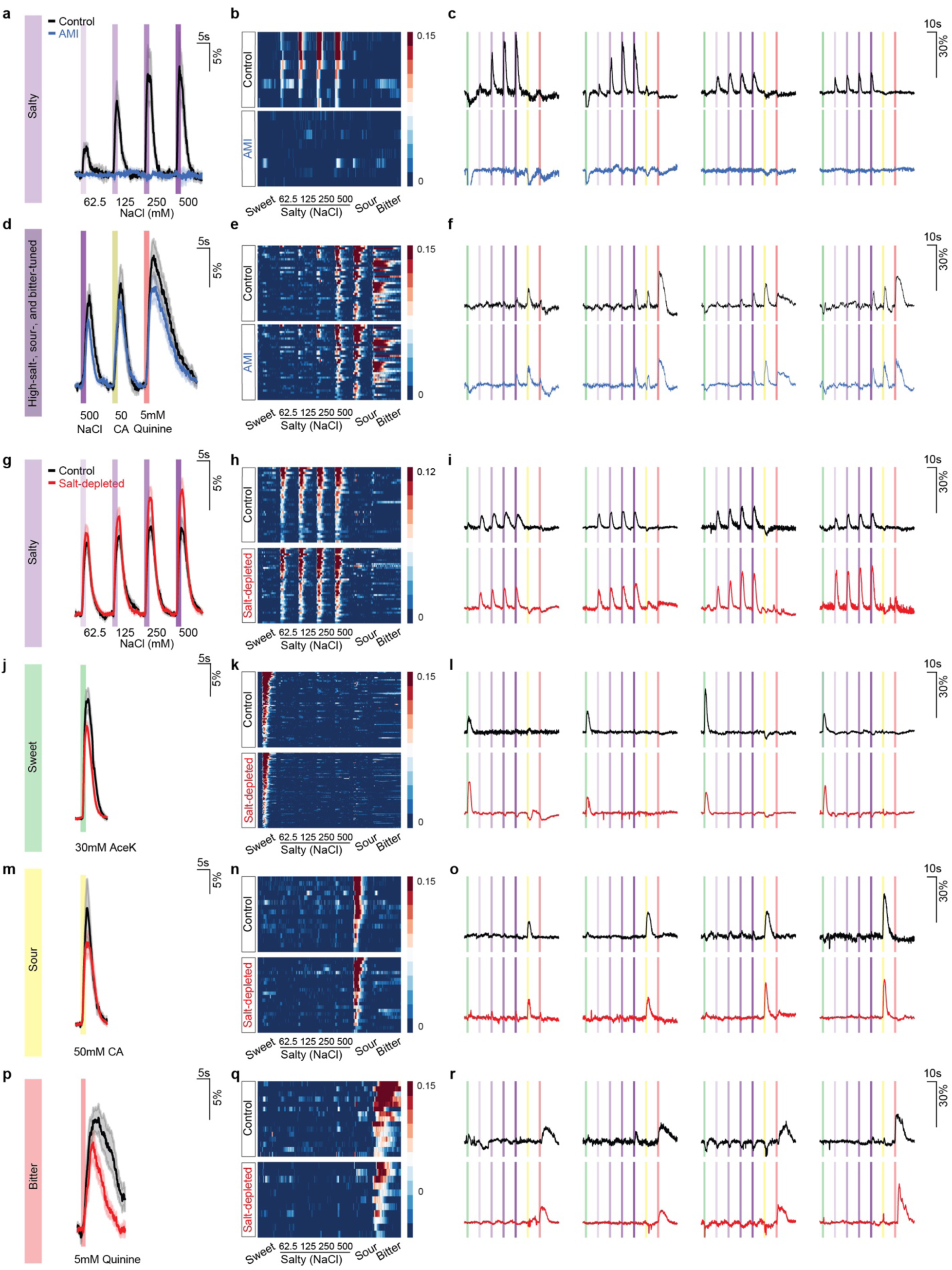
Salt-selective ganglion neurons show enhanced activity towards sodium salt after salt depletion. **a-f**. Amiloride (AMI) treatment eliminates responses from salt-tuned neurons (**a–c**) but not from bitter- and/or sour-tuned neurons that exhibit sensitivity to high salt (**d–f**; labeled as high-salt-, sour-, and bitter-tuned). **a,d.** Average traces of salt-selective (**a**, *n* = 8) or high-salt-, sour-, and bitter-tuned (**d**, *n* = 35) neurons in control animals before (black) and after (blue) amiloride treatment. Shaded areas indicate s.e.m.; bars represent 2-s stimulation windows, with stimuli listed at the bottom. Scale, dF/F. **b.e.** Heatmaps depicting fluorescence traces from geniculate ganglion neurons identified as salt-tuned (**b**) or high-salt-, sour-, and bitter-tuned (**e**) neurons. Stimuli are listed at the bottom. **c, f.** Representative traces from four neurons in each category. Stimuli are 2-s oral infusions of sweet (30 mM acesulfame potassium, AceK), salt (62.5, 125, 250, and 500 mM NaCl), sour (50 mM citric acid), and bitter (5 mM quinine) solutions. **g-r.** Calcium imaging responses of geniculate ganglion neurons in control (Ctrl, n=451 from 19 mice) and salt-depleted (SD, n=720 from 25 mice) animals. **g,j,m,p.** Average unnormalized traces of salt-selective neurons (Ctrl, n = 31; SD, n = 39) in response to 62.5, 125, 250, and 500 mM NaCl (**g**), sweet-selective neurons (Ctrl, n = 68; SD, n = 106) in response to 30 mM AceK (**j**), sour-selective neurons (Ctrl, n = 16; SD, n = 22) in response to 50 mM citric acid (**m**), and bitter-selective neurons (Ctrl, n = 15; SD, n = 11) in response to 5 mM quinine (**p**). The corresponding normalized traces are shown in Fig. 1h–k. **h,k,n,q.** Heatmaps depicting fluorescence traces from geniculate ganglion neurons identified as salt-(**h**), sweet-(**k**), sour-(**n**), and bitter-selective neurons (**q**). **i,l,o,r.** Representative raw traces of 4 each of salty-(**i**), sweet-(**l**), sour-(**o**), or bitter-(**r**) selective neurons from control (black, top row) or salt-depleted mice (red, bottom row). Bars represent 2s windows of stimulations described in **c, f.**

**Extended data Fig. 3.**
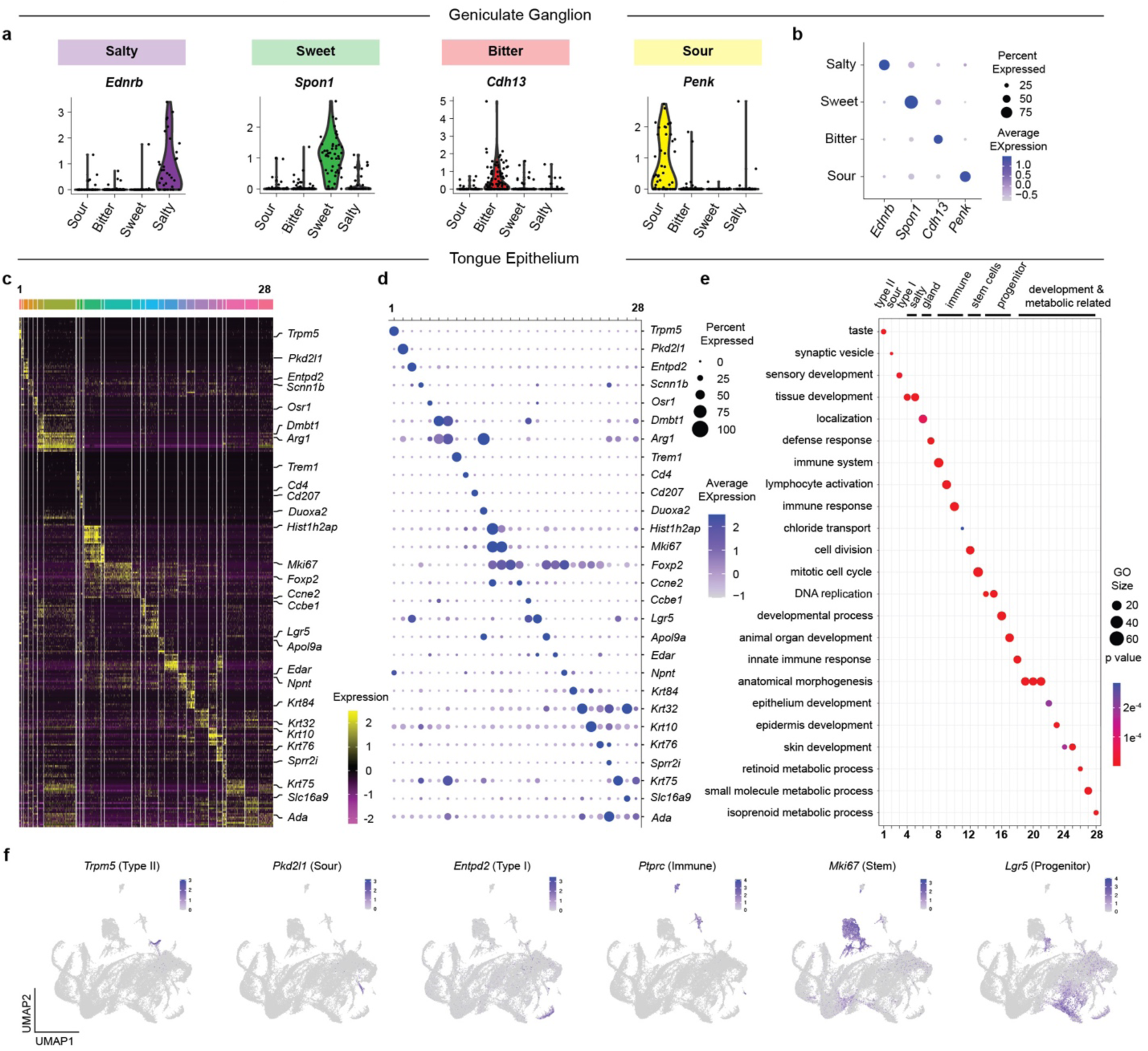
Single cell sequencing analyses of the geniculate ganglion neurons and tongue epithelial cells. **a-b**. Violin plots (**a**) and a dot plot (**b**) show marker genes for populations of geniculate ganglion neurons tuned to different tastes, analyzed using the dataset published in Zhang, et al., *Cell*, 2019. *Ednrb* is specifically expressed in salty-, *Spon1* in sweet-, *Cdh13* in bitter-, and *Penk* in sour-selective neurons. **c-d.** Heatmap illustrating expression of 10 gene markers for each cluster 1 to 28 (the list of genes is included in Extended Data Table 1). Individual clusters are separated by space. Expression levels are pseudo-colored based on z scores (purple to yellow). One representative marker gene from each cluster was labelled (**c**), and the same marker genes are plotted in the dot plot (**d**). **e.** Top 1 Gene Ontology term (abbreviated) per cluster for 28 cell clusters. Full lists of the top 10 GO terms per cluster, calculated using the top 100 expressed genes, are provided in Extended Data Table 1. **f.** Feature plots of gene markers that label 6 distinctive populations in the tongue—Type II taste cells (*Trpm5*), Type III sour-sensing taste cells (*Pkd2l1*), Type I glia-like cells (*Entpd2*), Putative immune cells (*Ptprc*), proliferating stem cells (*Mki67*), and lingual progenitors (*Lgr5*).

**Extended data Fig. 4.**
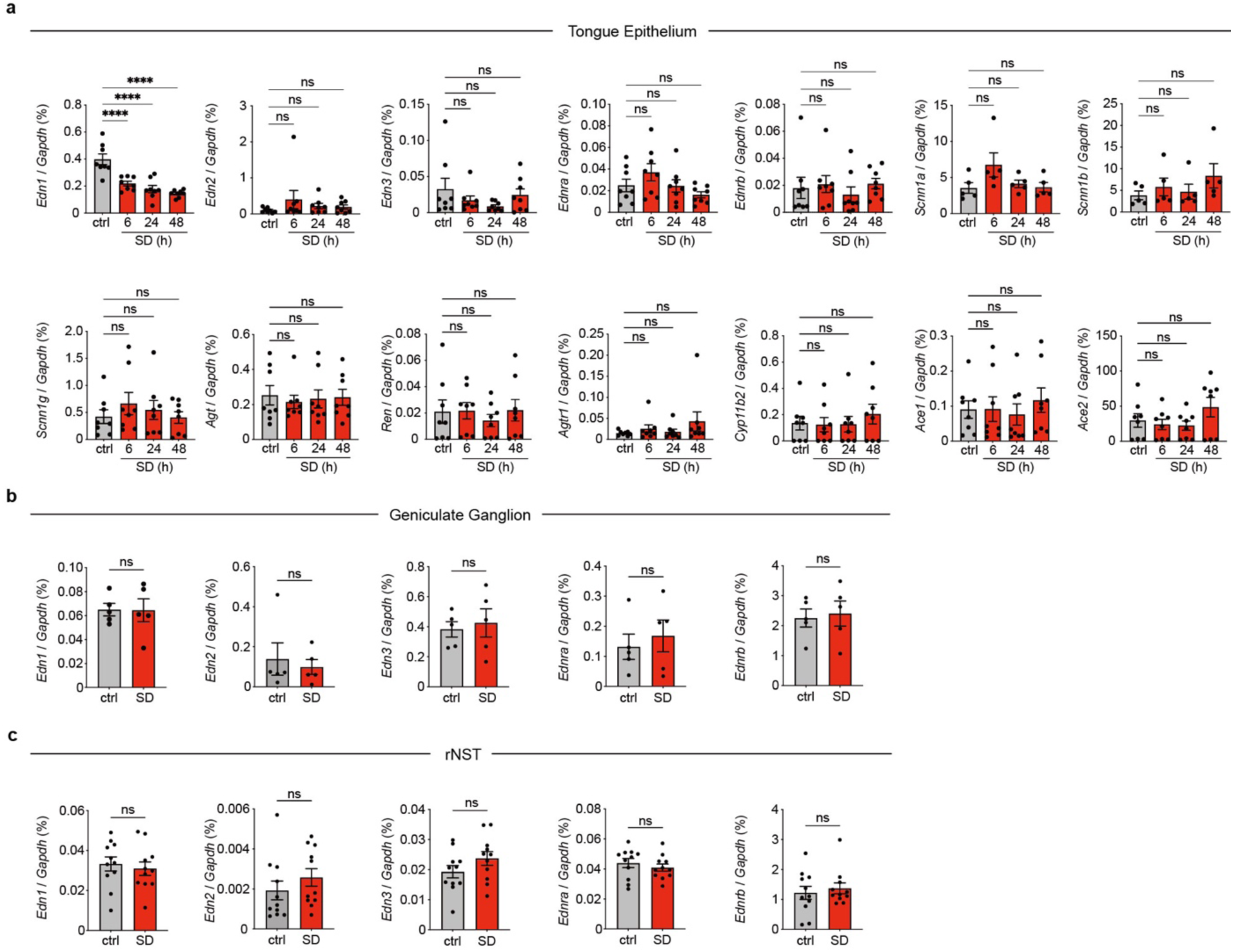
Expression of *Edn1* in the tongue epithelium gradually decreases after salt depletion. **a.** Quantitative RT-PCR (RT-qPCR) analysis of *Edn1*, *Edn2*, *Edn3*, *Ednra*, *Ednrb*, *Scnn1a* (encoding ENaCα), *Scnn1b* (encoding ENaCβ), *Scnn1g* (encoding ENaCγ), *Agt*, *Ren* (encoding Renin), *Agtr1*, *Cyp11b2*, *Ace1*, *Ace2*. Expression levels are normalized to *Gapdh*. *n* = 8, one-way ANOVA, Dunnett’s multiple comparisons test, *P* < 0.0001 (*Edn1*), *P* = 0.4741 (*Edn2*), *P* = 0.3198 (*Edn3*), *P* = 0.1191 (*Ednra*), *P* = 0.7753 (*Ednrb*), *P* = 0.1193 (*Scnn1a*), *P* = 0.4346 (*Scnn1b*), *P* = 0.6365 (*Scnn1g*), *P* = 0.9528 (*Agt*), *P* = 0.8480 (*Ren*), *P* = 0.4319 (*Agtr1*), *P* = 0.7645 (*Cyp11b2*), *P* = 0.8324 (*Ace1*), *P* = 0.2364 (*Ace2*). **b.** RT-qPCR analysis of *Edn1*, *Edn2*, *Edn3*, *Ednra*, and *Ednrb* in the geniculate ganglion, as normalized to *Gapdh*. Each dot represents 10 geniculate ganglia combined from 5 animals, *n* = 25 animals each group. Unpaired t-test, *P* = 0.9644 (*Edn1*), *P* = 0.6678 (*Edn2*), *P* = 0.6964 (*Edn3*), *P* = 0.6090 (*Ednra*), *P* = 0.7804 (*Ednrb*). **c.** RT-qPCR analysis of *Edn1*, *Edn2*, *Edn3*, *Ednra*, and *Ednrb* in the rostral nucleus of the solitary tract (rNST), as normalized to *Gapdh*. Each dot represents 1 animal, *n* = 11. Unpaired t-test, *P* = 0.6373 (*Edn1*), *P* = 0.3271 (*Edn2*), *P* = 0.1648 (*Edn3*), *P* = 0.4402 (*Ednra*), *P* = 0.5987(*Ednrb*).

**Extended data Fig. 5.**
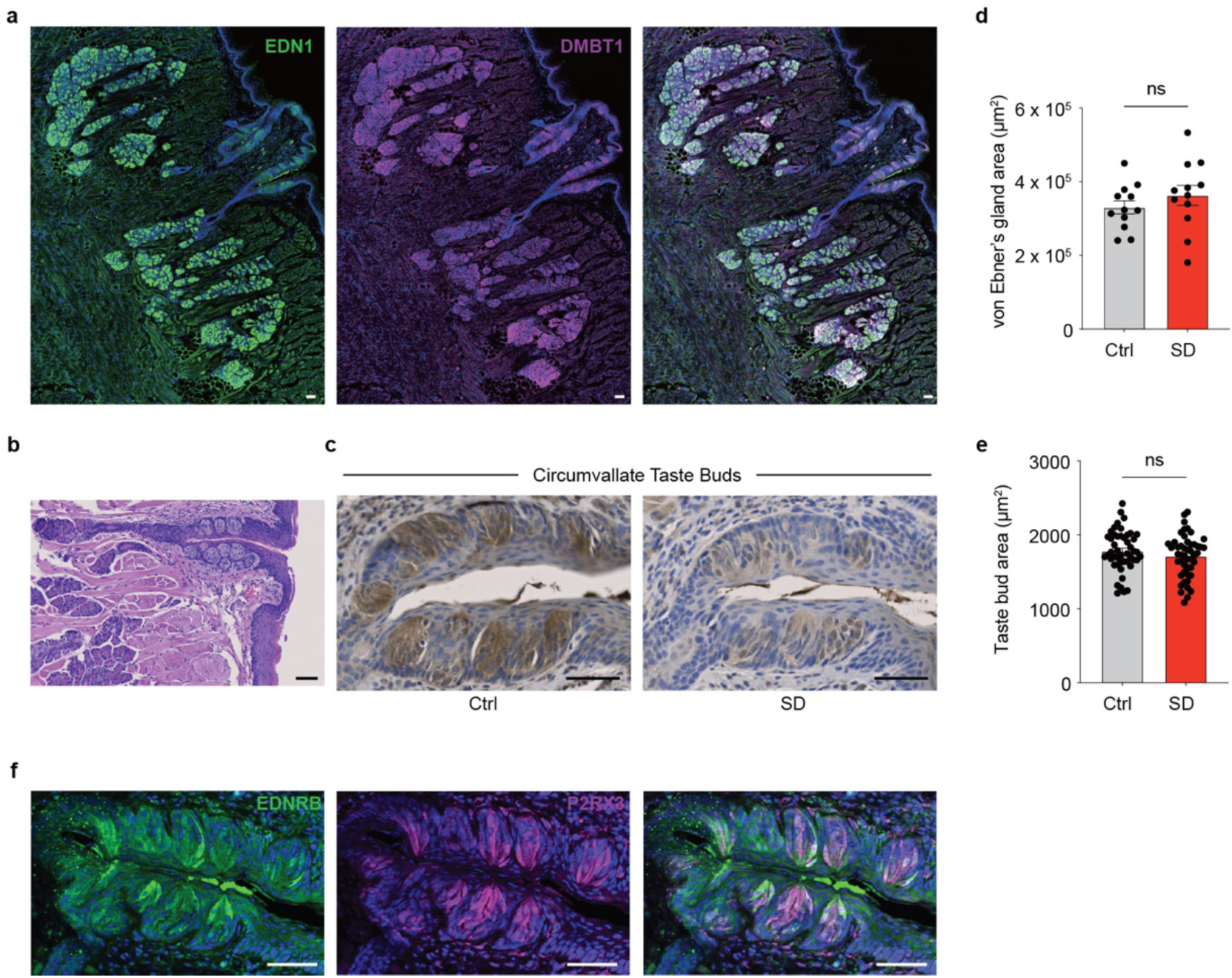
EDN1 is enriched in the tongue glands and taste buds. **a.** EDN1 (green) immunostaining shows that it is enriched in the von Ebner’s glands and the circumvallate papillae taste buds, colocalizing with DMBT1 (magenta). *n* = 6, a representative picture is shown. Left, EDN1 (green); middle, DMBT1 (magenta); right, composite. **b.** Haematoxylin-eosin staining showing the ducts of von Ebner’s glands connect to taste papillae (*n* = 3). **c.** Endothelin 1 (EDN1, brown) immunohistochemistry with haematoxylin staining in the circumvallate papillae taste buds of a representative control or salt-depleted animal (*n* = 4 each). **d, e.** Size quantification of von Ebner’s glands (**d**, *n* = 12) and taste buds (**e**, *n* = 48) in control (gray) and salt-depleted (SD, red) animals. **f.** Immunofluorescent staining showing EDNRB (green) and P2RX3 (magenta) at the nerve terminals of ganglion neurons innervating the circumvallate papillae taste buds. *n* = 4, a representative picture is shown. DAPI (blue). All scale bars, 50 µm.

**Extended data Fig. 6.**
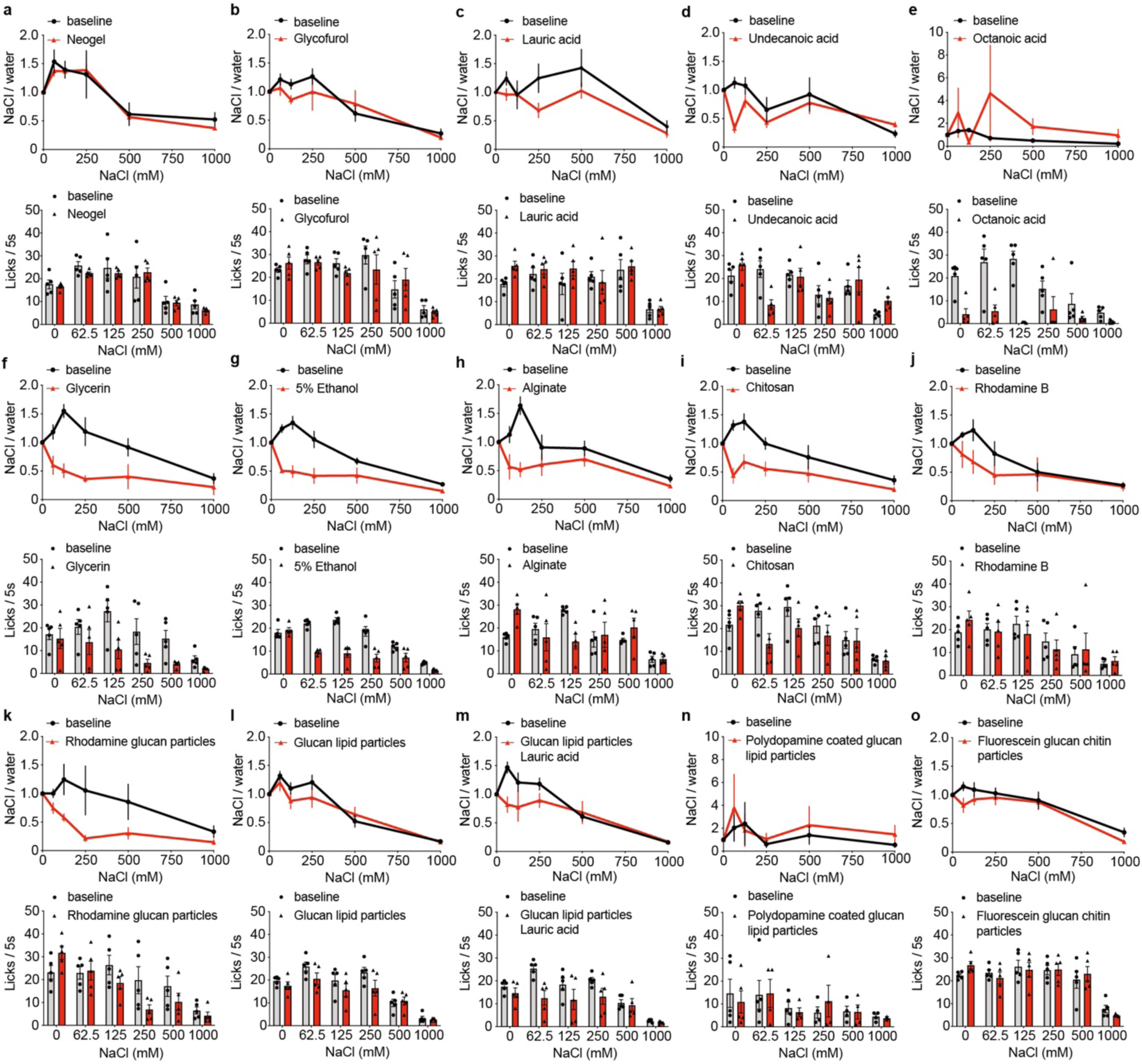
Characterization of compounds for specific tongue delivery. **a-o**, Top panel, lick ratio of NaCl to water after applying the candidate compounds on the tongue surface; bottom panel, the corresponding licks per 5s trial to water, 62.5, 125, 250, 500, and 1,000 mM NaCl. Two-way ANOVA, Šídák’s multiple comparisons test, *n* = 5, *P* = 0.7430 (**a**), *P* = 0.4413 (**b**), *P* = 0.0467 (**c**), *P* = 0.0921 (**d**), *P* = 0.4183 (**e**), *P* = 0.0030 (**f**), *P* = 0.0003 (**g**), *P* = 0.0165 (**h**), *P* = 0.0048 (**i**), *P* = 0.0963 (**j**), *P* = 0.0403 (**k**), *P* = 0.3148 (**l**), *P* = 0.1184 (**m**), *P* = 0.7380 (**n**), *P* = 0.0652 (**o**).

**Extended data Fig. 7.**
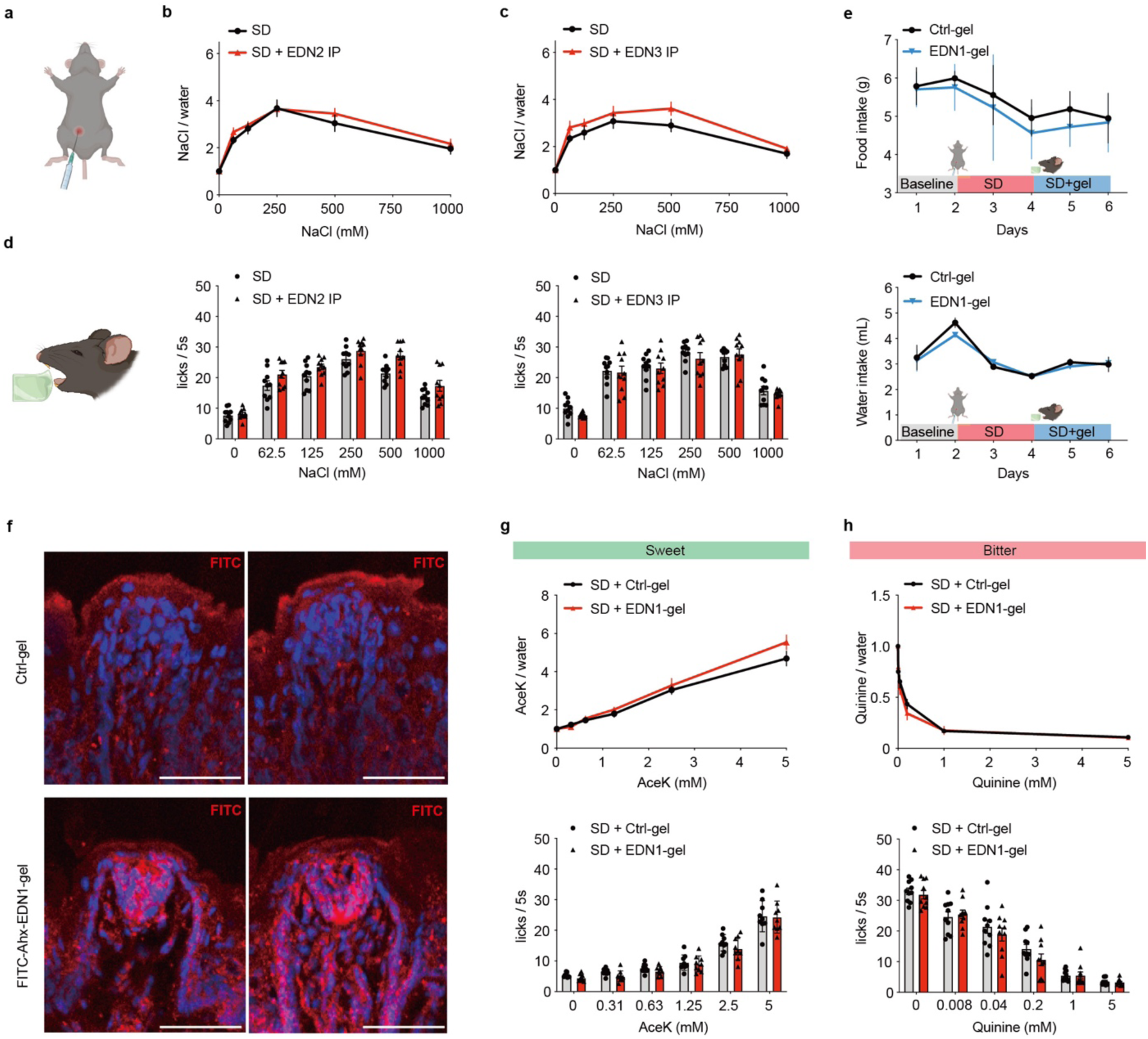
EDN2 and EDN3 do not affect salt taste, and EDN1 tongue applications do not alter sweet or bitter taste. **a.** Schematic illustration of intraperitoneal (i.p.) injection. **b,c**. Top, animals’ lick ratio of NaCl to water after i.p. injections of EDN2 (b) or EDN3 (c). Bottom, the corresponding licks per 5s trial to water, 62.5, 125, 250, 500, and 1,000 mM NaCl after EDN2 or EDN3 i.p. *n* = 10, Two-way ANOVA, Šídák’s multiple comparisons test, *P* = 0.4445 (b), *P* = 0.1451 (c). **d**. Schematic illustration of Neogel application to the tongue. **e**. Food (top) and water (bottom) intake before (baseline), during salt depletion (SD), and after application of Neogel (SD + gel) containing solvent (Ctrl-gel, black, *n* = 5) or EDN1 (EDN1-gel, blue, *n* = 4). **f**. Neogel containing FITC-Ahx-EDN1 was applied to the tongue (EDN1-gel, *n* = 6 mice), with delivery detected by anti-FITC immunostaining (red), and compared with empty Neogel (Ctrl-gel, *n* = 4 mice). Two representative fungiform taste buds are shown for each condition. **g-h**. Top, animals’ lick ratio of sweet (AceK, g) or bitter (quinine, h) to water after applications of Neogel containing EDN1. Bottom, the corresponding licks per 5s trial of water and sweet (g) or bitter (h) solutions. Two-way ANOVA, Šídák’s multiple comparisons test, *P* = 0.3012 (g, *n* = 9), *P* = 0.6774 (h, *n* = 10). Schematics in **a,b,e** were created using BioRender (http://biorender.com).

**Extended data Fig. 8.**
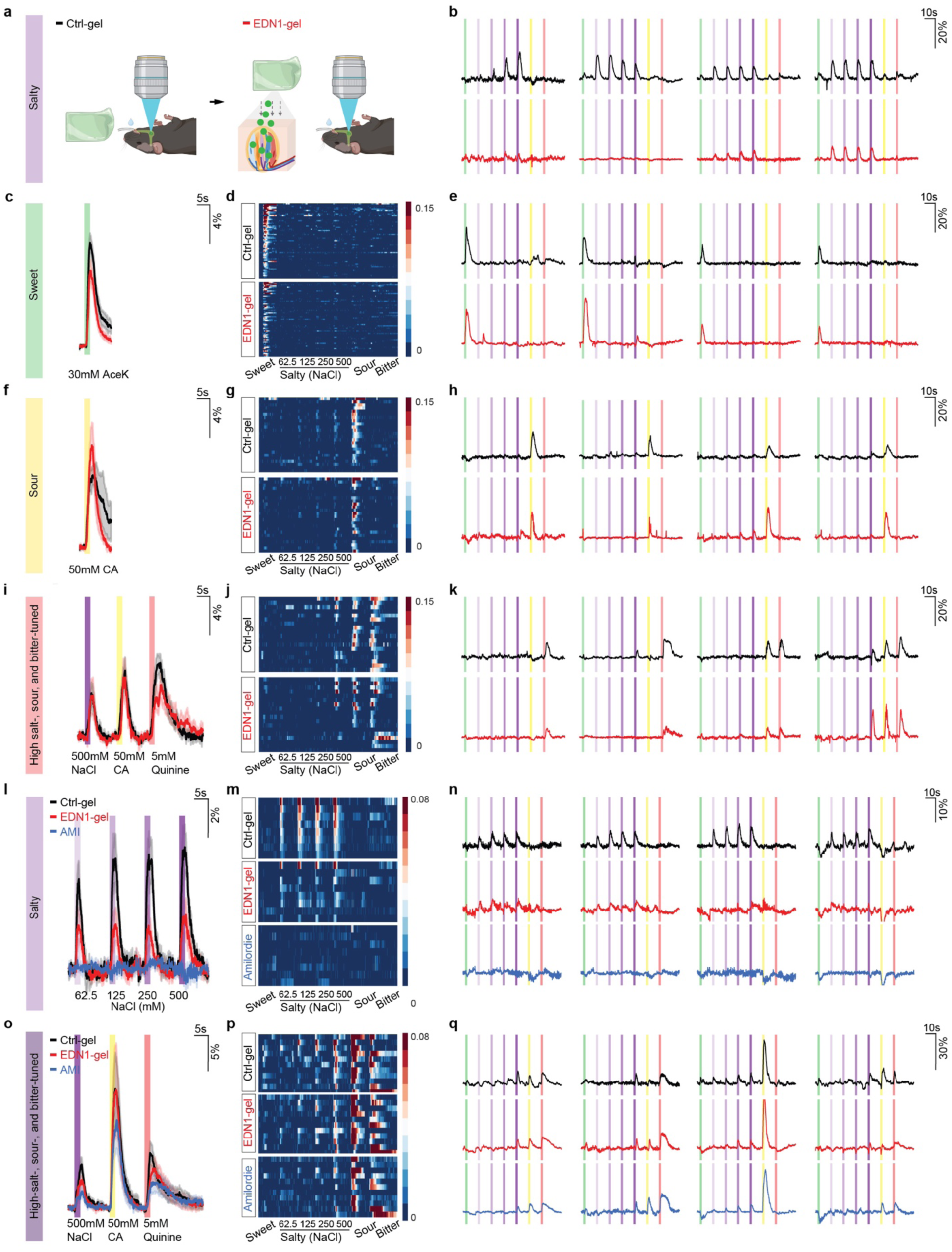
Neogel containing EDN1 specifically reduces the activity of salty-tuned neurons. **a.** Neogel containing solvent (Ctrl) or EDN1 was applied to the tongue of 48 h salt-depleted animals (**a-k**, *n* = 14 mice). Neuronal activity in response to tastant stimulation was recorded 30 min after each application. Responses of salt-selective neurons (*n* = 22) are shown in Fig. 3i–j. **b,e,h,k.**Tastant-triggered responses from 4 representative neurons tuned to salty (**b**), sweet (**e**), sour (**h**), or bitter (**k**) solutions after application of Ctrl-gel (black), or EDN1-gel (red). Bars represent 2s stimulation windows of sweet (30mM AceK, green), salty (62.5, 125, 250, and 500mM NaCl, light to dark purple), sour (50mM citric acid, yellow), and bitter (5mM quinine, red) solutions. Scale, dF/F. **c,f,i.** Average traces of sweet-selective neurons responding to 30mM AceK (**c,** *n* = 59), sour-selective neurons responding to 50mM citric acid (**f,** *n* = 24), and bitter-responding neurons (**i**, *n* = 18: 3 responded to bitter only, 6 to bitter and sour; 2 to bitter and 500mM NaCl, and 7 to bitter, sour, and 500mM NaCl. **d,g,j.** Heatmap of neuronal activity in response to tastant stimulations from sweet-selective (**d**), sour-selective(**g**), and bitter-responding neurons (**j**). Stimulations are listed at the bottom. **l-q.** Amiloride (AMI) treatment eliminates the responses of salt-tuned neurons (**l–n**) that are inhibited by EDN1-gel treatment in 48 h salt-depleted animals, but not those of bitter- and/or sour-tuned neurons that respond to high concentrations of NaCl (**o–q**; labeled as high-salt-, sour-, and bitter-tuned). Empty Neogel (Ctrl-gel) was first applied to the tongue, and neuronal responses were recorded, followed by application of EDN1-gel and recording of responses. AMI was then applied to confirm amiloride sensitivity. *n* = 7 mice. **l,o.** Average traces of salt-selective (**l**, *n* = 8) or high-salt-, sour-, and bitter-tuned (**o**, *n* = 11) neurons in control animals following application of Ctrl-gel (black), EDN1-gel (red), and amiloride (blue). Shaded areas indicate s.e.m.; bars represent 2-s stimulation windows, with stimuli listed at the bottom. Scale, dF/F. **m,p.** Heatmaps depicting fluorescence traces from geniculate ganglion neurons identified as salt-tuned (**m**) or high-salt-, sour-, and bitter-tuned (**p**) neurons following treatment with Ctrl-gel (top), EDN1-gel (middle), or amiloride (bottom). Stimuli are listed at the bottom. **n,q.** Representative traces from four neurons in each category. Stimuli were delivered as 2-s oral infusions, as listed in **m, p**. Schematics in **a** were created using BioRender (http://biorender.com).

**Extended data Fig. 9.**
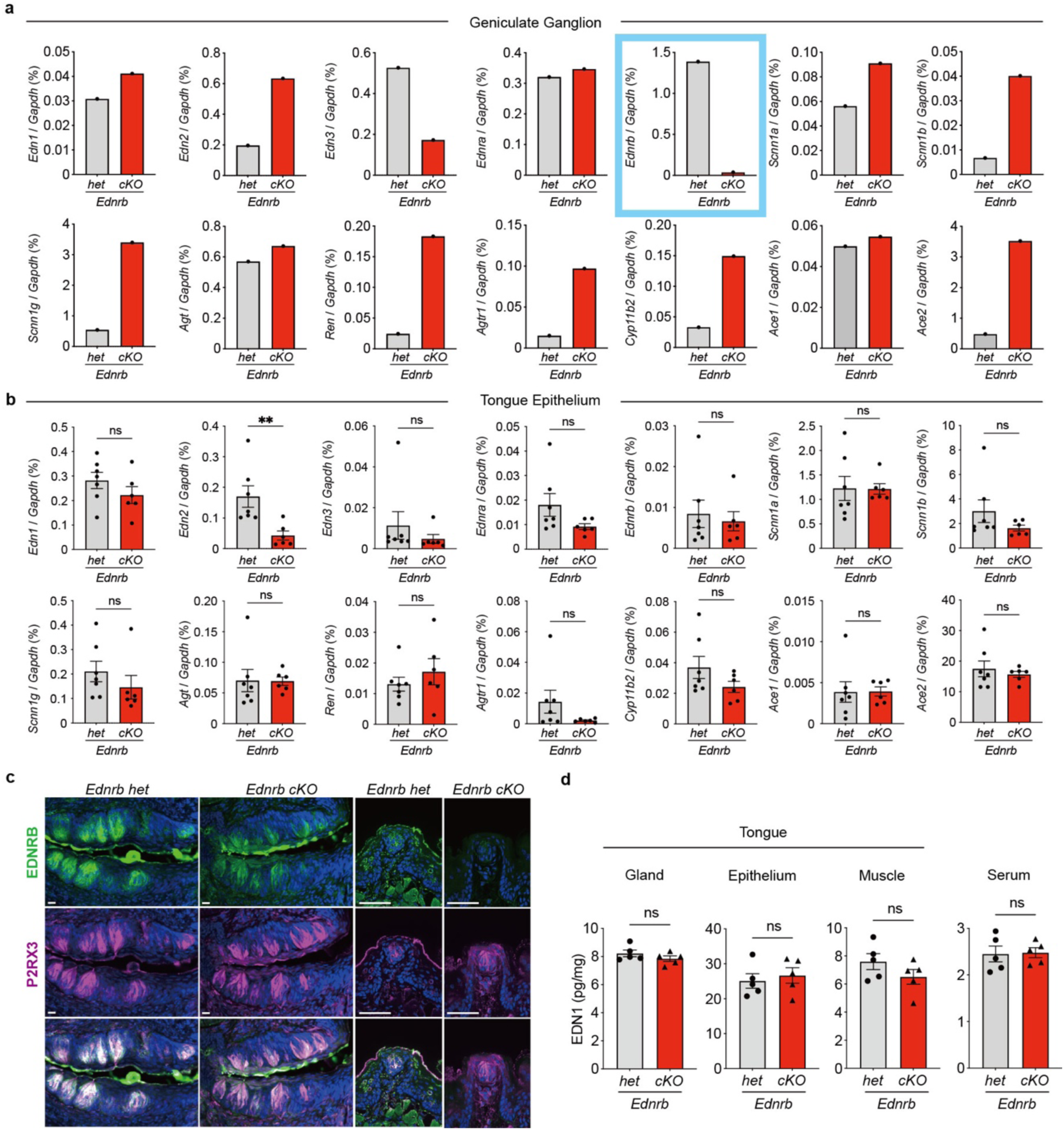
*Ednrb* conditional knockout animals show reduced mRNA in geniculate ganglia and protein at tongue projections. **a, b**. RT-qPCR analysis of *Edn1*, *Edn2*, *Edn3*, *Ednra*, *Ednrb*, *Scnn1a*, *Scnn1b*, *Scnn1g*, *Agt*, *Ren*, *Agtr1*, *Cyp11b2*, *Ace1*, and *Ace2* in the geniculate ganglion (a) and tongue epithelium (b), as normalized to *Gapdh*. Each dot represents 12 geniculate ganglia combined from 6 animals in a; *n* = 7 *Ednrb het* and *n* = 6 *cKO* animals in b. A blue box highlights the validation of diminished *Ednrb* mRNA levels in the geniculate ganglion neurons. b, unpaired t-test, *P* = 0.2386 (*Edn1*), *P* = 0.0095 (*Edn2*), *P* = 0.4066 (*Edn3*), *P* = 0.1131 (*Ednra*), *P* = 0.6783 (*Ednrb*), *P* = 0.9662 (*Scnn1a*), *P* = 0.2076 (*Scnn1b*), *P* = 0.3351 (*Scnn1g*), *P* = 0.9623 (*Agt*), *P* = 0.4002 (*Ren*), *P* = 0.1619 (*Agtr1*), *P* = 0.1646 (*Cyp11b2*), *P* = 0.9674 (*Ace1*), *P* = 0.5512 (*Ace2*). **c**. Immunostaining of EDNRB (green) and P2RX3 (magenta) in the circumvallate and fungiform taste buds of *Ednrb het* (*P2rx3-Cre; Ednrb^+/f^*) and *cKO* (*P2rx3-Cre; Ednrb^f/f^*) animals. *n* = 4 each group, a representative image is shown. Scale bars, 50µm. **d**. EDN1 ELISA shows no differences in the tongue von Ebner’s glands (Gland), epithelium, muscle, or the serum of *Ednrb het* and *cKO* animals. *n* = 5 each, unpaired t-test, *P* = 0.2617 (gland), *P* = 0.6213 (epithelium), *P* = 0.1999 (muscle), *P* = 0.8955 (serum).

**Extended data Fig. 10.**
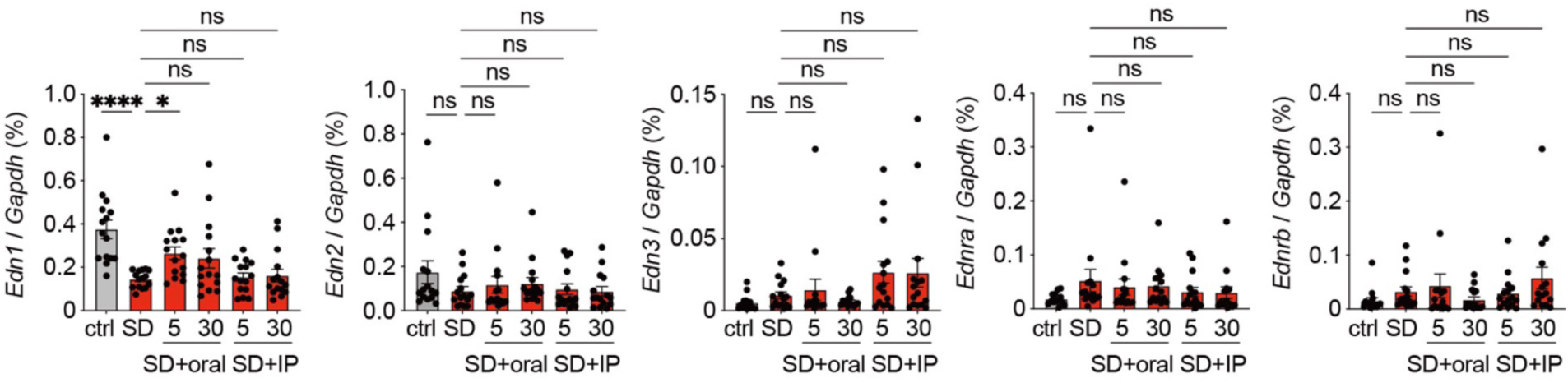
Oral NaCl delivery induces *Edn1* expression in 5 minutes. RT-qPCR analysis of *Edn1*, *Edn2*, *Edn3*, *Ednra*, and *Ednrb* in control (Ctrl), salt-depleted (SD), and SD animals 5 or 30min after oral licking or intraperitoneal (i.p.) injections of 1M NaCl. *n* = 10, One-way ANOVA, Dunnett’s multiple comparisons test, *Edn1*, compare to Ctrl, *P* < 0.0001 (SD), compare to SD, *P* = 0.0378 (SD+5min oral), *P* = 0.1307 (SD+30min oral), *P* = 0.9997 (SD+5min i.p.), *P* = 0.9954 (SD+30min i.p.); *Edn2*, compare to Ctrl, *P* = 0.2243 (SD), compare to SD, *P* = 0.9569 (SD+5min oral), *P* = 0.9088 (SD+30min oral), *P* = 0.9997 (SD+5min i.p.), *P* > 0.9999 (SD+30min i.p.); *Edn3*, compare to Ctrl, *P* = 0.9626 (SD), compare to SD, *P* = 0.9888 (SD+5min oral), *P* = 0.9875 (SD+30min oral), *P* = 0.2203 (SD+5min i.p.), *P* = 0.2430 (SD+30min i.p.); *Ednra*, compare to Ctrl, *P* = 0.2037 (SD), compare to SD, *P* = 0.9419 (SD+5min oral), *P* = 0.9747 (SD+30min oral), *P* = 0.6557 (SD+5min i.p.), *P* = 0.5964 (SD+30min i.p.); *Ednrb*, compare to Ctrl, *P* = 0.8504 (SD), compare to SD, *P* = 0.9696 (SD+5min oral), *P* = 0.8942 (SD+30min oral), *P* = 0.9997 (SD+5min i.p.), *P* = 0.5486 (SD+30min i.p.).

